# Decreased endothelial cell retinoic acid signaling accelerates progression of single ventricle pulmonary arteriovenous malformations

**DOI:** 10.64898/2025.12.08.693095

**Authors:** Henry Rousseau, Tina Wan, Nhi Nguyen, Jaime Wendt Andrae, Michael Tschannen, Angela J. Mathison, Victor Jin, Olivia Groh, Xingyan Zhou, Stryder M. Meadows, Ramani Ramchandran, Igor Shmarakov, Amy Y. Pan, Andrew D. Spearman

## Abstract

**Background:** Pulmonary arteriovenous malformations (PAVMs) are vascular complications that universally develop in patients with single ventricle congenital heart disease after Glenn surgery. However, the pathophysiological mechanisms underlying single ventricle PAVMs are poorly understood. To comprehensively evaluate molecular changes post-Glenn, we performed single-cell RNA sequencing (scRNAseq) on rat lung samples after Glenn surgery.

**Methods:** Using adult Sprague Dawley rats, we performed scRNAseq on unfiltered lung samples 3 weeks after left-sided Glenn or sham surgery. We compared endothelial cell (EC) differentially expressed genes (DEGs) in our model to two mouse models of hereditary hemorrhagic telangiectasia (HHT), a hereditary condition characterized by visceral AVMs. Finally, we modified the vitamin A (Vit A) content of Glenn and sham rat diets and re-assessed PAVM shunting and EC transcriptional differences.

**Results:** Using scRNAseq (n=4 Glenn, n=4 sham), we identified 13 transcriptionally distinct lung cell clusters, including 3 EC clusters (general, capillary, lymphatic), with pronounced differences between Glenn and sham in the general EC cluster (∼17% of genes). Comparison to HHT mouse models confirmed overlap of ∼18% of DEGs, including identification of significantly downregulated genes involved in and regulated by all-*trans* retinoic acid (ATRA) signaling in all 3 models. Dietary deficiency of Vit A intake, a precursor of ATRA, caused increased PAVM shunting (p<0.01) that was mitigated with excess dietary Vit A intake. Lastly, EC-specific RNAseq identified Vit A diet-induced gene expression differences, including regulation of PI3K signaling.

**Conclusions:** In this study, we report the novel application of scRNAseq to study mechanisms underlying single ventricle PAVMs in a surgical rat model. We identified multiple dysregulated biological processes in rat lung ECs post-Glenn, including decreased ATRA signaling and conserved gene expression patterns with HHT. Dietary modification of Vit A intake altered post-Glenn shunting and represents a novel potential therapeutic strategy for single ventricle PAVMs and HHT AVMs.

## Introduction

Pulmonary arteriovenous malformations (PAVMs) are abnormal vascular connections between arteries and veins that universally develop in patients with single ventricle congenital heart disease (CHD) [1–5]. Single ventricle PAVMs develop in patients after undergoing surgical palliation with a Glenn procedure, whereby pulmonary blood flow comes exclusively from the superior vena cava and hepatic vein blood is excluded from pulmonary blood flow [6–7]. Post-Glenn PAVMs cause progressive intrapulmonary shunting that exacerbates pre-existing hypoxemia [8]. However, the pathophysiology of single ventricle PAVMs remains unknown, including the molecular drivers of PAVM pathogenesis, affected cell types, and similarities and differences with other clinical AVM conditions.

Previous studies used large and small animal models of unilateral Glenn circulation to investigate focused pathways potentially altered in single ventricle PAVMs [7–15]. One previous study with a rat Glenn model utilized an unbiased approach to identify affected signaling pathways post-Glenn [16]. Tipps et al performed an RNA microarray using whole lung lysate and identified multiple dysregulated vascular remodeling pathways at 8 months post-surgery (i.e., later stages of PAVM pathophysiology). To date, though, no studies have comprehensively investigated whole lung transcriptional changes post-Glenn at single-cell resolution or in the early stages of PAVM initiation post-Glenn. To identify the cellular and molecular drivers of single ventricle PAVMs using an unbiased approach, we performed the first single-cell RNA sequencing (scRNAseq) experiment in an animal model of Glenn circulation during the initiation phase of PAVMs post-Glenn. We identified that lung endothelial cells (ECs) are the primarily affected cell type in single ventricle PAVMs. Among multiple dysregulated EC signaling pathways, we identified molecular overlap with brain and lung AVMs in independent models of hereditary hemorrhagic telangiectasia (HHT), including decreased all-*trans* retinoic acid (ATRA) signaling among all three AVM models. Finally, in our Glenn model, we identified that decreased dietary vitamin A (Vit A) intake further decreases EC retinoid levels post-Glenn and accelerates the progression of PAVM shunting. Collectively, our data identified conserved signaling pathways in single ventricle and HHT AVM pathogenesis, including ATRA signaling as a novel potential therapeutic strategy.

## Methods

### Animals

Adult Sprague-Dawley rats (male and female, 6-8 weeks of age) were used for all experiments (Taconic). All rats were housed in the Biomedical Research Center at the Medical College of Wisconsin with access to standard chow diet, water ad libitum, and maintained in a 12-hour light/dark cycle. All rat surgical procedures described below were performed under isoflurane anesthesia (1-3%). All experimental rat protocols were approved by the Medical College of Wisconsin Institutional Animal Care and Use Committee prior to initiation of experimental protocols (Animal Use Agreement #7731).

*Smad4* EC-specific, tamoxifen-inducible knockout mice were generated as previously described [17] using conditional *Smad4*^f/f^ mice [18] and *Cdh5*-Cre^ERT2^ mice [19]. To induce *Smad4* gene deletion, 300 ug of tamoxifen (Sigma, T5648) was delivered orally at postnatal day (P)1, P2 and P3. *Smad4* mice were housed in ventilated cages within a temperature-controlled room with 12-hour light/dark cycle. *Smad4* animal protocols were approved and performed in accordance with Tulane University’s institutional Animal Care and Use Committee policies and adhere to ethical standards.

### Left-sided superior cavopulmonary anastomosis (Left-sided Glenn)

To model Glenn circulation in rats and create unilateral PAVMs, we performed an end-to-end unilateral cavopulmonary anastomosis between the left superior vena cava (L-SVC) and LPA, as previously described [15]. For a comparative control animal, a sham surgery was performed on an age and sex matched cage-mate, as previously described [15]. After completion of Glenn and sham surgeries, the chest was closed, isoflurane was stopped, and ventilation was continued until the rat demonstrated consistent spontaneous ventilation. Ventilation was then stopped, and the endotracheal tube was removed. Animals recovered in a clean and warmed cage until fully ambulatory. Upon complete recovery, Glenn and sham rats were returned to their housing in the Biomedical Research Center where they were co-housed until end-point experiments at 3 weeks post-surgery. This time point was chosen to evaluate early molecular changes after Glenn surgery during PAVM initiation, while avoiding immediate post-surgical changes.

### Rat lung dissociation & processing

Prior to tissue harvesting, rats were anesthetized with isoflurane and exsanguinated by abdominal aorta transection. The chest was then opened to expose the lungs and the left lung was gently perfused to clear blood using sterile PBS with a 25g needle in the L-SVC (Glenn) or main pulmonary artery (sham). The left lung was then harvested and divided in two below the level of the left main bronchus entering the left hilum. Each half of the left lung was then finely minced with scissors in separate sterile 3cm cell culture dishes. Minced tissue was then transferred to a gentleMACS C tube (Miltenyi, #130-096-334) for mechanical and chemical dissociation (multi tissue dissociation kit 2, Miltenyi, #130-110-203) using a gentleMACS OctoDissociator (Miltenyi, #130-096-427), per manufacturer recommendations. After removing the samples from the OctoDissociator, enzymatic digestion was neutralized by adding 7.5 ml of pre-warmed 10% FBS (ThermoFisher, #A5670401) in RPMI media (ThermoFisher, #11875093). The sample was then filtered through a pre-wet 30µm MACS smartstrainer (Miltenyi, #130-110-915) and centrifuged at 600 x *g* for 10 minutes. Supernatant was then aspirated and resuspended in 2 ml of freshly prepared RBC lysis buffer (ThermoFisher, #00-4300-54). After incubating for 3 minutes and gently pipetting with a wide bore pipette tip (ThermoFisher, #2079GPK), RBC lysis was quenched by adding 10 ml of 0.04% BSA (ThermoFisher, #AM2618) in sterile PBS. The sample was then centrifuged at 400 x *g* for 10 minutes. After aspirating the supernatant, the sample was resuspended in 0.04% BSA in PBS for scRNAseq or resuspended in MACS buffer (0.5% BSA, 2mM EDTA, and PBS; Miltenyi, #130-091-376 and #130-091-222) for cell sorting with magnetic microbeads or for flow cytometry. To minimize sampling bias, only the lower half of the left lung was used for scRNAseq. To optimize yields for RNA isolation and flow cytometry, both the lower and upper halves of the left lung were combined.

### scRNAseq library preparation and sequencing

To minimize potential technical differences between Glenn and sham samples, left lower lung samples from Glenn and sham rats were processed in pairs on four separate days (n=4 pairs, n=8 total) using 10x Chromium GEM-X single cell 3’ reagent kit. Cell counts and viability were evaluated in the Luna-FL automated counter (Logos Biosystems, Acridine Orange/Propidium Iodide Stain) with all samples exhibiting >92% viability. To further minimize potential technical differences, library preparation and next-generation sequencing were completed on all samples in one batch at Medical College of Wisconsin Mellowes Center for Genomic Science and Precision Medicine (RRID:SCR_022926). 10x Genomics Chromium workflow was initiated with approximately 10,000 cells per sample and processed according to the Chromium Next GEM Single Cell 3’ Reagent Kits (10X Genomics, sample dual indexes with cellular and molecular barcodes) using the 10X Genomics Chromium Controller. Fragment analysis (Agilent 5200 Fragment Analyzer System) was utilized to check the quality at cDNA and final library preparation stages, with 14 cycles of PCR used to amplify the single-cell libraries. Prior to sequencing, libraries were quantified and pooled by qPCR (Kapa Library Quantification Kit, Kapa Biosystems) and sequencing was completed on the Illumina NovaSeq 6000 targeting 50,000 reads per cell per condition. scRNAseq data were deposited in the Gene Expression Omnibus (GEO) for independent analysis (GSE312833).

### scRNAseq bioinformatic analysis data processing and quality control

scRNAseq raw reads were assigned to Ensembl Rat Genome (reference version Rnor_6.0.98) transcripts using the CellRanger (10X Genomic Cloud analysis) with default parameters. The processed data of each sample were then analyzed in R package Seurat (v5.0.0) [20–21]. For quality control, we excluded cells with unique feature counts of ≤200 or ≥2500, and cells with a mitochondrial gene fraction ≥5%. Global scaling was applied to normalize counts across all cells in each sample. Approximately 2,000 most variable genes in the dataset were used for dimensionality reduction by principal component analysis (PCA). Cell clusters were determined using FindNeighbours and FindClusters functions based on the first 17 PCs. Uniform Manifold Approximation Projection (UMAP) algorithm was used to visualize the cell clusters. Global gene expression distances were calculated using Euclidean distance of the average expression within each cell type, for each dataset [22–23]. The optimal number of cell clusters was determined using the Modularity Optimizer (version 1.3.0) and Louvain algorithm. To annotate cell clusters visualized in the combined UMAP, we performed Wilcoxon rank sum test to identify enriched gene expression within each cluster. Genes with average log2 fold change (log2FC) > 1 and expression in > 10% of cells within each cluster were compared with published references of rat, mouse, and human lung [23–30]. To compare the abundance of cells within each cluster, we quantified the number of cells within each cluster for each sample relative to the total cells for that sample.

### Differential gene expression between Glenn and sham rats

To identify differentially expressed genes (DEGs) between Glenn and sham rats within each cluster, model-based analysis of single-cell transcriptomics (MAST) was performed. We used cut-offs of log2FC >|0.5| and adjusted p-value <0.05 to identify DEGs within each cluster. To identify biological pathways dysregulated within each cluster, we uploaded DEGs into DAVID and entire cluster datasets (gene list, log2FC, and adjusted p-value) into Ingenuity Pathway Analysis (IPA).

### Sub-clustering of vascular endothelial cells

To identify potential sub-populations of affected vascular endothelial cells, we performed sub-clustering using all cells identified in clusters 1 and 3 at the previous step. After normalization and scaling, sub-clusters were determined using FindNeighbours and FindClusters functions based on the first 30 PCs. The optimal number of cell clusters was determined using the Modularity Optimizer (version 1.3.0) and Louvain algorithm. To annotate cell clusters visualized in the combined UMAP, we performed Wilcoxon rank sum test to identify enriched gene expression within each cluster and compared to published references for lung EC subtypes [23, 31–33].

### Rat lung endothelial cell isolation for scRNAseq validation

To validate scRNAseq data from lung ECs, we isolated CD31^+^ cells from left lung samples from independent Glenn and sham rats at 3 weeks post-surgery. After lung dissociation and resuspension in MACS buffer (0.5% BSA, 2mM EDTA, and PBS; Miltenyi, #130-091-376 and #130-091-222), we separated CD31^+^ lung cells via magnetic microbeads. First, we performed indirect negative selection to remove CD45^+^ lung cells (anti-rat CD45-FITC; BD, #554877; and anti-FITC microbeads; Miltenyi, #130-048-701) using MS columns (Miltenyi, #130-042-201). After collecting the CD45^-^ population, we performed indirect positive selection to capture CD31^+^ lung cells (anti-rat CD31-PE; BD, # 555027; and anti-PE microbeads; Miltenyi, #130-048-801) using MS columns. After isolation of CD45^-^CD31^+^ lung cells, we extracted RNA (PureLink RNA mini kit; ThermoFisher, #12183018A; with homogenizer columns; ThermoFisher, #12183026) for subsequent qPCR. Total RNA (250 ng) was transcribed into cDNA (iScript gDNA clear cDNA synthesis kit; BioRad, #1725035), and qPCR was performed (PR1MA qMax green qPCR mix; MidSci, #PR2000-N-1000) on the CFX96 qPCR machine (BioRad). Quantification was performed using the ΔΔCt method using *Gapdh* as the housekeeping gene. Primers for qPCR were previously published or custom-designed using Primer 3 (**Supp Table 1**) [34].

### Secondary analysis of endothelial bulk RNAseq

To identify potential molecular overlap with AVMs in HHT and single ventricle PAVMs, we compared our EC cluster 1 dataset to EC-specific RNAseq datasets of two independent HHT mouse models: a previously published Acvrl1/Alk1 brain AVM model [17] and Smad4 lung AVM model [35–36]. RNAseq data were obtained from isolated brain and lung ECs harvested from EC-specific, tamoxifen-inducible *Acvrl1* (*Alk1^iBEC^*) and *Smad4* (*Smad4^iLEC^*^)^ knock-out mice, respectively. Both models isolated brain and lung ECs using Dynabeads (Invitrogen 11035) coated with anti-CD31 antibody (BD Biosciences, #553370), as previously described [17,35]. Bulk RNAseq was subsequently performed as previously described using a NextSeq 550 System (Illumina, SY-415-1002) [37].

The differential expression analysis was performed on raw gene counts using DESeq2 (v1.47.5) after pre-filtering to include only genes that have a count of at least 10 for half of the samples [38]. DEGs were identified in both HHT AVM datasets by adjusted p-value <0.05 and then compared to DEGs previously identified in our EC cluster 1 Glenn dataset (Glenn DEGs: adjusted p-value <0.05 and log2FC >|0.5|). After identifying DEGs common to all 3 datasets, we identified overlapping DEGs that were also differentially expressed in the same direction in all 3 datasets (i.e., concordant). RNAseq data for re-analyzed public datasets are found in GEO (GSE197105), and RNAseq data for newly analyzed datasets were deposited in GEO for independent analysis (GSE312375).

### Dietary modification of Vitamin A intake

Based on identification of shared dysregulation of ATRA-related genes, we sought to determine whether further decreasing or increasing lung EC retinoid levels could impact PAVM initiation and/or progression by modifying dietary intake of Vit A. Glenn and sham rats were started on custom diets (Teklad Diets, Inotiv) 1-2 weeks prior to surgery and continued for 3 weeks after surgery. Vit A control diet contained 20 IU/g Vit A, Vit A deficient diet contained 0 IU/g Vit A, and Vit A excess diet contained 200 IU/g Vit A (**Supp Table 2**). Custom diets were identical except for Vit A content. Tissue (lung, liver, and plasma) retinoid levels (retinol [ROH] and retinyl esters [RE]) were analyzed to assess the Vit A status of animals on different diets. Additionally, the diets contained 610 ppm aspirin with an estimated daily dosing of 5-10 mg/kg/day, which is equivalent to pediatric clinical dosing of aspirin for children with Glenn circulation.

### Quantification of intrapulmonary shunting

To assess the functional impact of PAVMs, we objectively quantified intrapulmonary shunting within the left lung, as previously described [15]. Briefly, a beveled catheter (MRE-027, Braintree Scientific) was directly inserted into the distal left pulmonary artery and secured by suture. The lungs were then re-inflated and ventilated. The left atrium was opened to allow unrestricted pulmonary vein egress, and the left lung was perfused with 10 ml of PBS. We then injected 10µm fluorescent microspheres (ThermoFisher, F8834) (20 µl microspheres diluted in 500 µl PBS) immediately followed by 1 ml PBS flush. Microspheres and PBS flush were directly aspirated from the left atrium using a wide bore pipette tip (ThermoFisher, 2079GPK). Microspheres were then aliquoted into a 96-well plate and quantified using a fluorescent plate reader (SpectraMax i3x, ex/em 580/605). Shunt fraction was calculated as a percentage relative to the fluorescence of non-injected microspheres: (collected spheres/non-injected spheres) x 100.

### Flow cytometry quantification of intracellular retinoid levels

To quantify the prevalence of retinoid-storing cells and intracellular retinoid levels, we utilized spectral cytometry based on previously characterized endogenous retinoid auto-fluorescence [39–41] at emission wavelength 455nm. After lung dissociation, we stained freshly isolated lung samples with anti-CD31-PE antibody (BD Biosciences, #555027) and viability dye (Live/Dead Yellow, Thermo, #L64968). Samples were incubated at 1:100 antibody concentration in MACS buffer for 20 minutes at 4^°^C, washed, and filtered immediately prior to running samples. Using a Cytek Spectral Cytometer (Cytek Aurora) without autofluorescence extraction, we quantified the median fluorescent intensity (MFI) of all live cells at UV5 channel as a read-out for the amount of intracellular retinoids (ROH and REs). We also quantified the abundance of retinoid^+^ cells, MFI at UV5 channel within retinoid^+^ cells, abundance of retinoid^+^CD31^+^ cells, and MFI at UV5 channel within retinoid^+^CD31^+^ cells.

### HPLC quantification of tissue retinoid levels

High-performance liquid chromatography (HPLC) was utilized to quantify tissue retinoid levels from tissue samples collected from post-surgical rats. Solid tissue samples were collected from the upper segment of the left lung and the lateral aspect of the left lobe of the liver. Solid tissue samples were immediately flash frozen in liquid nitrogen. Whole blood was collected from the right external jugular vein of anesthetized rats. Blood was placed into K2 EDTA tubes (Braintree Scientific, MV-500 201339-BX), immediately centrifuged at 2000 x *g* for 15 minutes, and then plasma was aliquoted and frozen at -80°C.

ROH and REs were extracted from plasma, liver, and lung under a dim yellow light and analyzed by HPLC, as previously described [40]. Briefly, livers and lungs were homogenized in 10 vol of PBS (10 mM sodium phosphate, pH 7.2, 150 mM sodium chloride) using a tissue tearer. An aliquot of plasma (100 µL) or tissue homogenate (equivalent to 50 µg of tissue) was then treated with an equal volume of absolute ethanol containing a known amount of retinyl acetate as an internal standard. The retinoids present in either plasma or tissue homogenates were extracted into hexane. The hexane extract was dried under a stream of N2 and redissolved in a running solvent. For determination of tissue and plasma levels of ROH and REs, a reverse-phase HPLC method employing a 4.6 × 250 mm Symmetry C18 Column (Waters, Milford, MA, USA) and a running solvent consisting of acetonitrile/methanol/methylene chloride (70:15:15 v/v) flowing at 1.8 mL/min was used. An Agilent 1260 Infinity II HPLC system (including Agilent 1260 Infinity II Binary Pump, Agilent 1260 Infinity II Vialsampler, Agilent 1260 Infinity II Ultrasensitive Diode Array Detector, and Agilent 1260 Infinity II Multicolumn Thermostat) controlled by Agilent OpenLab ChemStation version 2.3.54 software was employed for these analyses. ROH and REs were detected at 325 nm. Quantitation was based on comparisons of the area under the peaks and spectra for unknown samples to those of known amounts of standards. The recovery of the internal standard was employed to correct for loss during extraction. For this analysis, HPLC-grade solvents were purchased from ThermoFisher Scientific. Quantified retinoid levels (ng of either ROH or total REs per sample) were normalized to DNA levels (µg of DNA per sample) and expressed as ng/µg DNA. The DNA concentration of each sample was assessed with a High Sensitivity Quant-iT™ dsDNA Assay Kit (ThermoFisher Scientific) according to the manufacturer’s protocol.

### Bulk RNAseq analysis of sorted CD31^+^ lung cells

To determine how dietary modification of Vit A intake impacted lung EC gene expression after Glenn surgery, we performed bulk RNAseq on microbead-sorted CD31^+^ lung cells 3 weeks after Glenn or sham surgery. After isolating CD31^+^ lung cells from post-surgical rats, we extracted RNA (PureLink RNA mini kit; ThermoFisher, #12183018A; with homogenizer columns; ThermoFisher, #12183026) for subsequent RNAseq. Library preparation was completed at Medical College of Wisconsin Mellowes Center for Genomic Science and Precision Medicine, utilizing the SMART-Seq Ultra Low v4 Kit (Takara). Briefly, RNA was reverse transcribed into cDNA and polyA enriched before Illumina adapters and indexes were added by PCR (Illumina Nextera XT DNA library prep). Sequencing was subsequently performed on the NovaSeqX platform at University of Wisconsin-Madison Center for Genomic Science Innovation targeting 40 million unique reads per sample. RNAseq output was aligned to the Norway rat genome (mRatBN7.2/rn7). RNAseq data were deposited in GEO for independent analysis (GSE311758).

Differential gene expression was analyzed using DESeq2 (v1.47.5), as previously described [38]. We first identified DEGs (adjusted p-value <0.05) between Glenn and sham rats on each diet. Next, we compared DEGs among all 3 diets to identify overlapping and unique DEGs. From the unique DEGs on each diet, we identified impacted biological pathways using DAVID.

### Statistical Analysis

Data are expressed as mean and standard deviation (SD) for continuous data and n (%) for categorical data unless otherwise stated. Bioinformatic approaches for scRNAseq and bulk RNAseq analyses are detailed above. We used unpaired t-tests with multiple comparison adjustment (Benjamini two-stage step-up) to compare relative cell counts within each cluster between Glenn and sham rats. We used Wilcoxon signed rank test to evaluate gene expression differences with qPCR. Differences among group in retinoid levels were analyzed using a t test or ANOVA, with the false discovery rate (FDR) applied to adjust for multiple comparisons. Where necessary for parametric assumptions, log transformation was employed. A p-value < 0.05 was considered statistically significant unless otherwise specified. Analyses were performed using GraphPad Prism 10 (GraphPad Software, San Diego, CA) and SAS 9.4 (SAS Institute Inc., Cary, NC).

## Results

### scRNAseq dataset of rat lungs after Glenn or sham surgery

To identify gene expression differences during PAVM initiation stages after Glenn surgery, we performed scRNAseq in Glenn and sham rats 3 weeks after surgery using the 10X Genomics system (**Fig 1A**). With 8 samples (n=4 Glenn and n=4 sham), we initially included a total of 79,846 cells (average of 9,980 cells/sample). After applying conservative QC filters, we included a total of 26,454 cells (average of 3,307 cells/sample), which included 15,388 cells (58.2%) from Glenn rats and 11,066 cells (41.8%) from sham rats.

**Figure 1.**
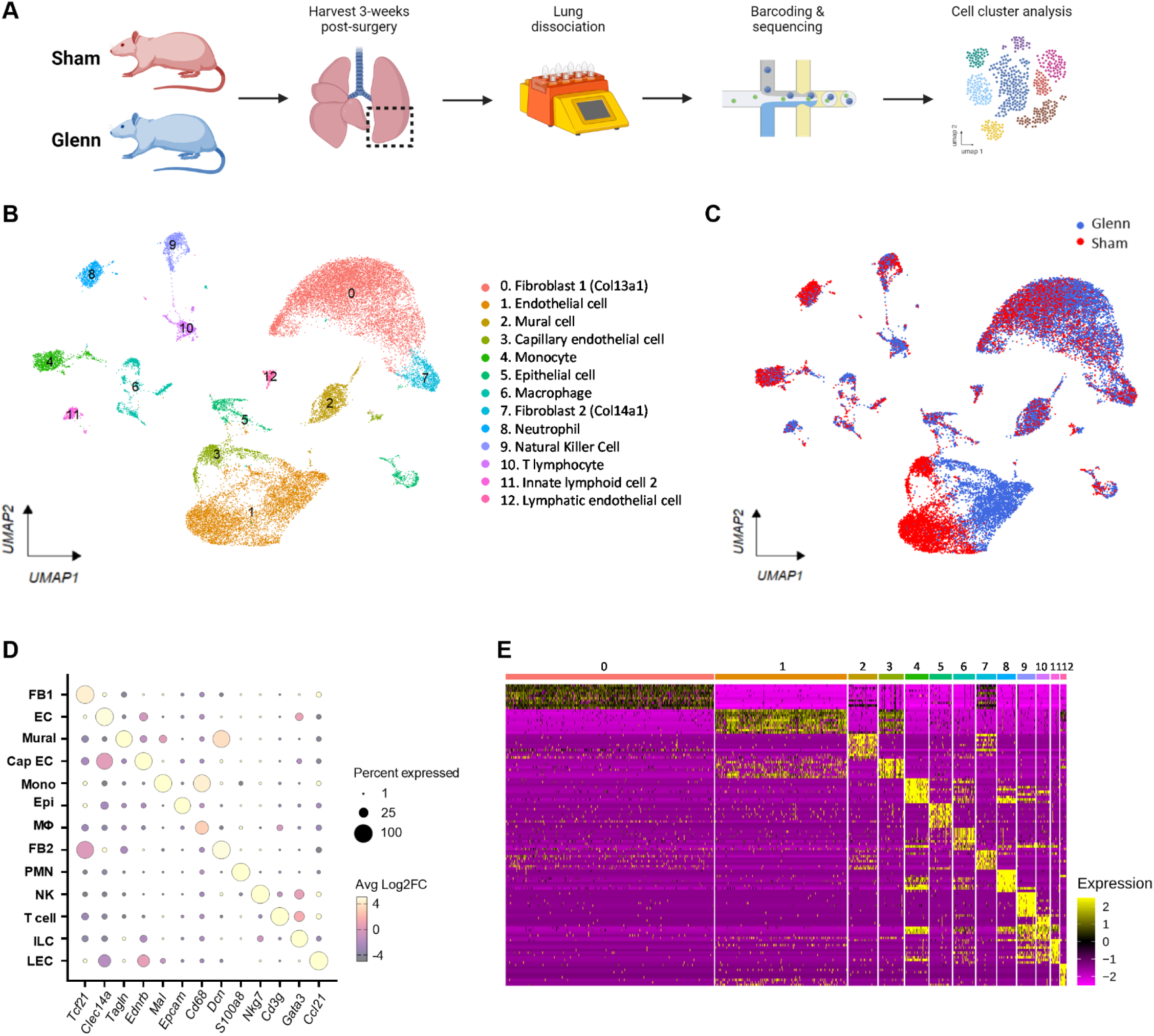
Single-cell RNA sequencing (scRNAseq) in rats early after Glenn or sham surgery identifies distinct clusters of lung cells. (A) Schematic for experimental workflow for performing scRNAseq in Glenn (n=4) and sham (n=4) rats. (B) Combined uniform manifold approximation and projection (UMAP) plot with all samples (n=8) showing distinct cellular clusters identified with unsupervised clustering and subsequent cluster annotation. (C) UMAP plot color coded by surgical group (Glenn versus sham) demonstrates pronounced separation within cluster 1 (general lung endothelial cells) and cluster 3 (capillary lung endothelial cells) between Glenn and sham rats. (D) Dot plot of marker genes identifying each cluster. Dot size indicates percent of cells within each cluster expressing the specific gene and dot color indicates the average log2 fold change of gene expression within each cluster compared to all other clusters. (E) Global heatmap of all clusters showing transcriptional gene expression differences for the top 5 genes within each cluster. Each column represents an individual cell and each row represents a gene.

### Altered lung cell populations

Based on unsupervised cell clustering of our dataset and annotation based on available lung references [23–33], we identified 13 distinct cell clusters, including 3 lung EC clusters (**Fig 1B-E**). To visualize global transcriptional differences between Glenn and sham groups within clusters, we color coded the UMAP by surgical groups. This UMAP demonstrated transcriptional variation within clusters that was most pronounced within cluster 1, which is the general EC cluster (**Fig 1C**). Importantly, Glenn/sham harvest pairs (**Supp Fig 1A**) and individual samples (**Supp Fig 1B**) were distributed across all clusters, suggesting no evidence of technical bias or artifact.

To quantitatively compare gene expression differences between Glenn and sham rats in all clusters, we quantified the abundance of DEGs within each cluster (**Fig 2A**). Cluster 1 showed the most discrepant transcriptional signatures with 16.7% (1310/7849) of all genes differentially expressed between Glenn and sham rats. Cluster 0 and cluster 3 also showed marked differences with 4.9% (355/7215) and 3.7% (321/8303) of genes differentially expressed, respectively. All other clusters had ≤ 1.3% genes differentially expressed. Next, we quantified whole transcriptome gene expression differences within each cluster using a Euclidean distance-based statistical approach [22–23], which validated that cluster 1 ECs were the most transcriptionally distinct cell type between Glenn and sham rats followed by cluster 0 fibroblasts and cluster 3 capillary ECs (**Fig 2B**). Last, to identify potential differences in cell numbers between surgical groups in each cluster, we compared the relative abundance of cells within each cluster. There were significantly fewer lung capillary ECs (cluster 3) in Glenn rats compared to sham rats (Glenn 2.7±0.4%, sham 6.9±1.6%; q-value =0.0252); however, there were no significant differences in cell numbers in any other cluster (**Fig 2C**). Altogether, these data indicate that ECs are the primarily affected cell type in the lung early after Glenn palliation, including that there is potential loss of capillary ECs as early as 3 weeks after Glenn.

**Figure 2.**
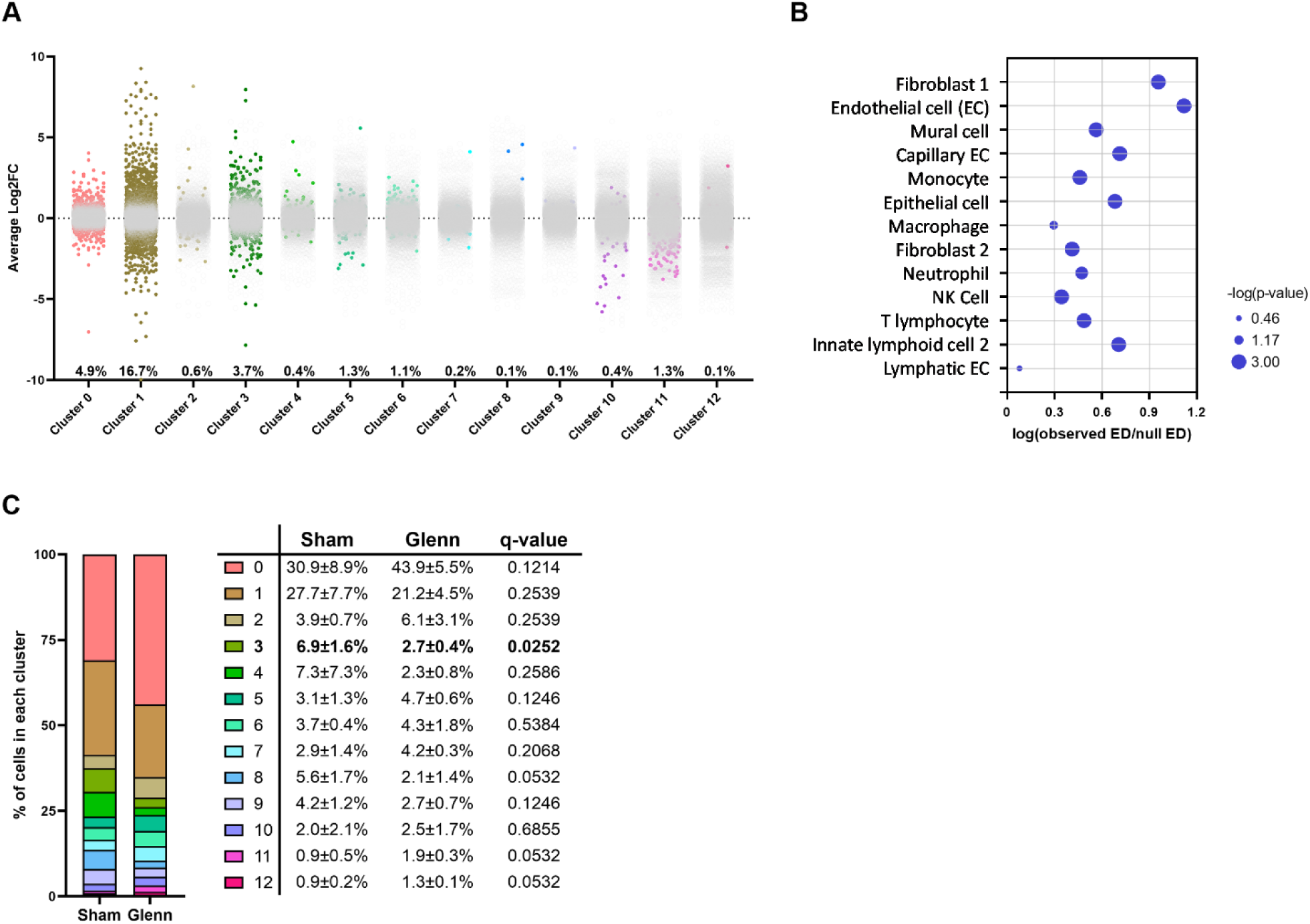
Lung endothelial cells (ECs) are the predominantly affected cell type early after Glenn surgery. (A) Scatter plot of all clusters showing all genes identified within each cluster. Colored dots indicate differentially expressed genes (DEGs), defined as average log2FC >|0.5| and adjusted p-value <0.05. Gray dots indicate genes within each cluster that were not differentially expressed. Percent annotated along the x-axis indicates the percent of genes within each cluster that were differentially expressed between Glenn and sham. Cluster 1 ECs were again the most strongly affected cluster with 16.7% of all genes differentially expressed. (B) Whole transcriptome comparison of global gene expression differences between Glenn and sham rats within each cluster based on the Euclidean-distance based statistical approach. Cluster 1 ECs had the most distinct gene expression differences between Glenn and sham. (C) Stacked bar graph with accompanying table indicates the mean ± SD of percent cells within each cluster from each rat. Q-value indicates the adjusted p-value for multiple unpaired t-tests comparing the frequency of cells within each cluster. Cluster 3 capillary ECs were significantly decreased in Glenn rats compared to sham rats; no differences between groups in all other clusters.

### Differentially expressed genes in rat Glenn lung endothelial cells

Based on the pronounced differences visualized by the UMAP plot (**Fig 1C**) and quantified in **Fig 2** suggesting that ECs are the primarily affected cell type in the lung early after Glenn palliation, we identified and compared DEGs within EC clusters 1 and 3 between Glenn and sham rats.

Within cluster 1 ECs, we identified 755 (57.6%) up-regulated genes (average Log2FC >0.5, adjusted p-value <0.05) and 555 (42.4%) down-regulated genes (average Log2FC <-0.5, adjusted p-value <0.05) (**Fig 3A**). Using all DEGs within cluster 1 (n=1310), we used DAVID pathway analysis to identify dysregulated biological processes (**Fig 3B**). Angiogenesis was the most significantly dysregulated pathway (p=1.9E-13). Multiple pathways overlapping angiogenesis and vascular remodeling included positive regulation of cell migration (p=8.7E-8), positive regulation of cell population proliferation (p=8.9E-8), and vasculogenesis (p=7.9E-5). Additionally, several unexpected inflammatory response pathways were identified, including response to LPS (p=1.4E-10), cellular response to TNF (p=1.4E-6), cellular response to IL-1 (p=8.6E-6), and response to cytokine (p=3.1E-5). Next, to further investigate dysregulated signaling pathways with a second bioinformatic tool and quantify directionality of altered signaling, we used IPA to identify dysregulated canonical signaling pathways (**Fig 3D**). Multiple canonical pathways supported the DAVID analysis identifying angiogenesis, including increased phosphatidylinositol-3 kinase (PI3K)/AKT signaling (Z-score +2.6, p=2.5E-3), increased TGF-β signaling (Z-score +2.3, p=8.9E-6), decreased angiopoietin signaling (Z-score -1.3, p=8.3E-4), and decreased bone morphogenetic protein (BMP) signaling (Z-score -2.3, p=1.1E-8). We also re-identified pathways associated with inflammation, including pathogen induced cytokine storm (Z-score +2.6, p=3.5E-4) and macrophage migration inhibitory factor (MIF) regulation of innate immunity (Z-score +2.4, p=1.3E-4). Finally, we identified decreased activity of key regulatory pathways, including signaling by retinoic acid (Z-score -2.0, p=2.7E-2)], antioxidant action of vitamin C [Z-score -1.9, p=8.3E-4], and transport of vitamins [Z-score -1.3, p=4.2E-3]. Altogether, cluster 1 DEGs and altered signaling pathways support that early PAVM initiation is likely multi-factorial and may be driven by pro-angiogenic/pro-inflammatory stimuli and derangement of regulatory pathways controlled by retinoic acid and vitamin C.

**Figure 3.**
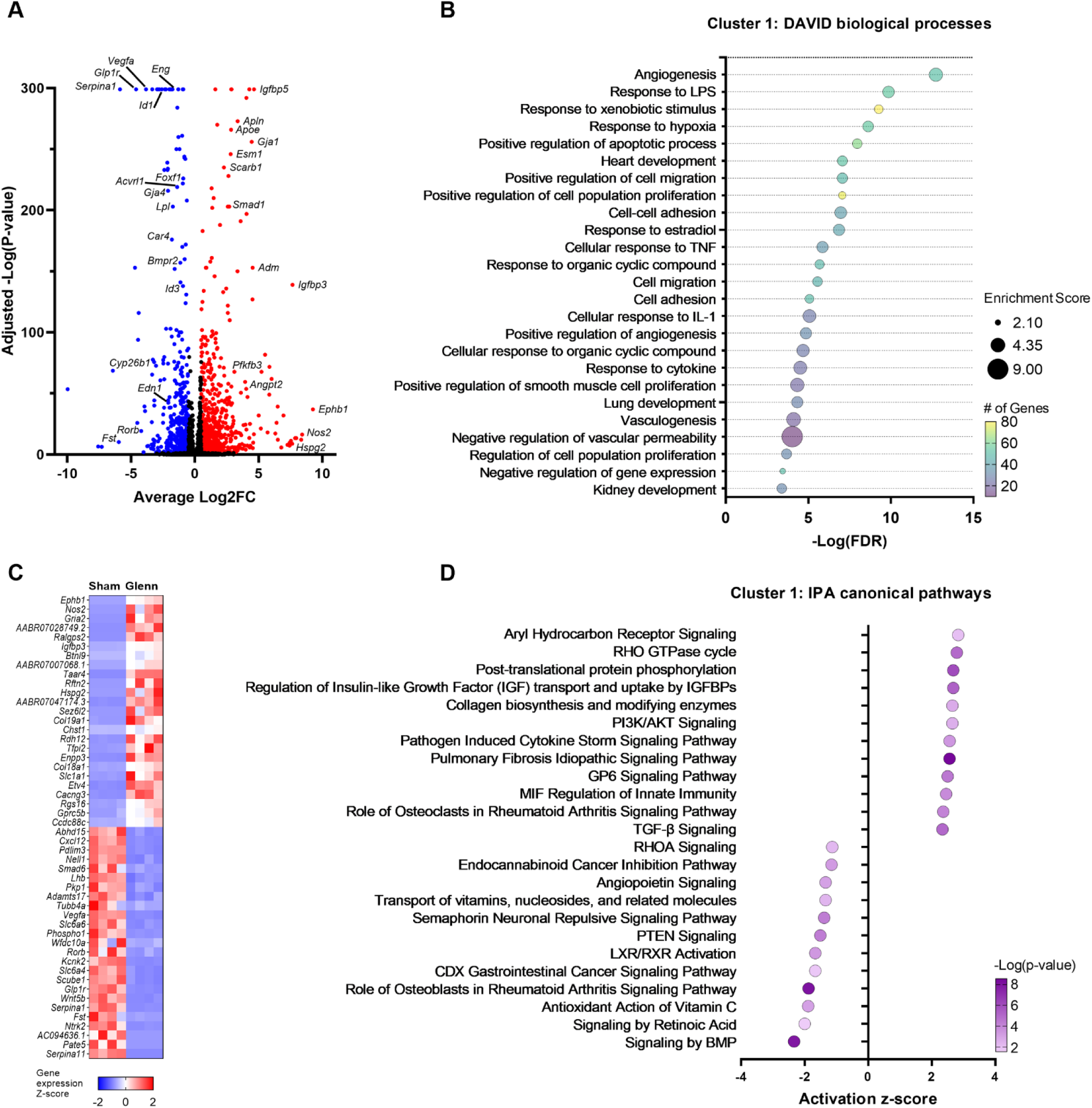
General lung endothelial cells (ECs) comprise cluster 1 cells and are significantly dysregulated in Glenn versus sham rats. (A) Volcano plot showing upregulated (red) and downregulated (blue) genes identified in cluster 1 with annotation of notable DEGs. (B) Dot plot showing the top 25 biological pathways identified using DAVID pathway analysis and cluster 1 DEGs. (C) Heatmap of the top 25 up-regulated and top 25 down-regulated DEGs in cluster 1. Each column represents an individual rat, and each row represents a gene. (D) Dot plot of the 12 most activated and 12 most de-activated canonical pathways identified using Ingenuity Pathway Analysis (IPA) and the entire cluster 1 dataset. Positive activation z-score indicates increased activity of the associated pathway in Glenn rats, whereas negative activation z-score indicates decreased activity of the associated pathway in Glenn rats.

Within cluster 3, we identified 165 (51.4%) genes up-regulated (average Log2FC >0.5, adjusted p-value <0.05) and 156 (48.6%) genes down-regulated (average Log2FC <-0.5, adjusted p-value <0.05) (**Supp Fig 2A**). Using all DEGs within cluster 3 (n=321), we used DAVID pathway analysis to identify dysregulated biological processes (**Supp Fig 2B**). Similar to cluster 1, angiogenesis was the most significantly dysregulated pathway (p=1.4E-14) along with multiple other vascular remodeling pathways, including positive regulation of angiogenesis (p=6.6E-7), blood vessel remodeling (p=1.2E-5), vasculogenesis (p=4.6E-5), and more. Similarly, we again identified multiple inflammatory response pathways, including response to LPS (p=4.6E-7), cellular response to IL-1 (p=9.7E-6), cellular response to cytokine stimulus (p=4.8E-4), and response to cytokine (p=2.3E-3). Next, we again used IPA to independently identify dysregulated canonical signaling pathways (**Supp Fig 2D**). Similar to cluster 1, we identified angiogenic/vascular remodeling pathways (activin inhibin signaling pathway [Z-score +2.1, p=7.2E-5] and signaling by Notch3 [Z-score +2.0, p=3.5E-3]) and inflammatory pathways (IFNα/β signaling [Z-score +2.4, p=2.6E-4], pathogen induced cytokine storm [Z-score +1.9, p=8.3E-4], but decreased IL-1 signaling [Z-score -2.2, p=2.2E-2]).

We further sub-clustered vascular lung ECs (clusters 1 and 3) to identify whether Glenn circulation preferentially impacted arterial, venous, or capillary ECs. Based on unsupervised clustering, the two largest clusters in our sub-clustering analysis did not segregate based on classical arterial, venous, or capillary markers (**Supp Fig 3A-B**). Rather, the largest clusters (sub-clusters 0 and 1) differentiated based on previously identified DEGs between Glenn and sham ECs without evidence of technical bias based on Glenn/sham harvest pair or individual sample (**Supp Fig 3C-D**). Quantifying the contribution to each sub-cluster, there were significant differences (q-value < 0.0001) in the number of cells from each surgical group within sub-clusters 0 and 1 (**Supp Fig 3E**). We again identified fewer capillary ECs (sub-cluster 2) in the Glenn group (q-value = 0.0086). Global heatmap of EC sub-clusters identifies the top 10 enriched genes within each sub-cluster (**Supp Fig 3F**).

### Validation of scRNAseq endothelial gene expression with isolated rat lung endothelial cells

To validate scRNAseq DEGs, we performed qPCR from isolated CD31^+^ lung cells from independent Glenn (n=6) and sham (n=6) rats (**Fig 4; Supp Fig 4**). Multiple genes in angiogenic pathways were confirmed to have significantly different RNA levels that were consistent with our scRNAseq dataset, including increased *Angpt2* (p=0.02), *Cxcr4* (p<0.01), and *Esm1* (p<0.01), as well as decreased *Acvrl1* (p<0.01), *Tmem100* (p=0.04), and *Vegfa* (p<0.01). We also confirmed consistent gene expression differences in inflammation (increased *Il6st* [p<0.01], increased *Sparcl1* [p<0.01], decreased *Serpina1* [p<0.01]), metabolism (decreased *Lpl* [p<0.01], increased *Pfkfb3* [p<0.01]), and vasodilation (increased *Nos2* [p=0.03] and decreased *Edn1* [p<0.01]) (**Fig 4C**). Interestingly, we also confirmed decreased expression of multiple lung capillary markers, including both aerocyte (Cap1) capillary markers (*Car4,* p=0.01; *Ednrb*, p<0.01) and general lung (Cap2) capillary markers (*Plvap*, p<0.01; *Slc6a4*, p<0.01) (**Supp Fig 5**) [24].

**Figure 4.**
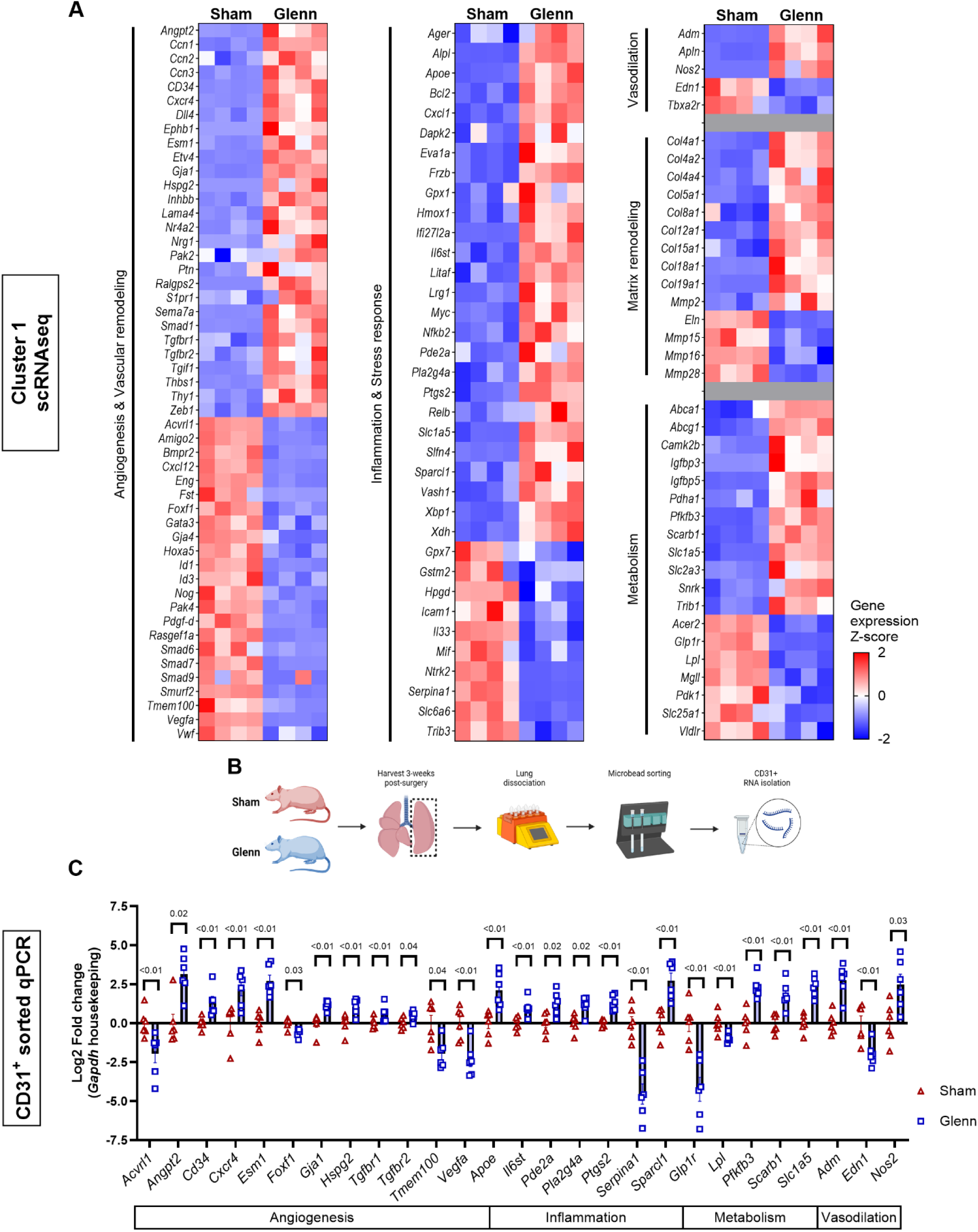
Glenn circulation impacts multiple signaling pathways in lung endothelial cells. (A) Heatmaps of gene expression z-scores for differentially expressed genes (DEGs) in cluster 1 endothelial cells organized by general endothelial cell function: angiogenesis/vascular remodeling, inflammation & stress response, vasodilation, matrix remodeling, and metabolism. (B) Schematic for experimental methodology for isolating RNA from CD31^+^ sorted lung cells from sham (n=6) and Glenn (n=6) rats. (C) Scatter plot with bars (mean ± SD) shows validation of specific gene targets using quantitative RT-PCR (qPCR). Genes are grouped by general biological processes, including angiogenesis, inflammation, metabolism, and vasodilation. Biological processes in (A) and (C) are not mutually exclusive as genes frequently perform multiple functions.

### Comparison of endothelial gene expression in HHT and post-Glenn AVMs

Based on gene expression patterns observed in post-Glenn lung ECs that recapitulated gene expression patterns observed in HHT pathophysiology (*Acvrl1, Angpt2, Esm1, Tmem100*, and more) [17,37,42–43], we hypothesized that there may be shared endothelial dysregulation in post-Glenn AVMs and HHT AVMs. To identify potential shared gene expression patterns, we compared our list of endothelial DEGs to two independent datasets derived from Smad4 and Acvrl1 HHT mouse models utilizing isolated ECs from lungs and brains, respectively (*Smad4*^iLECs^ and *Alk1*^iBECs^) (**Fig 5**) [17]. We identified a total of 755 and 4828 DEGs in the *Smad4* and *Alk1* datasets, respectively, in addition to the previously identified 1310 DEGs in cluster 1 of our rat Glenn scRNAseq dataset (**Fig 5A**). Comparing gene lists among all 3 datasets, we identified 136 (18.0%) overlapping DEGs and 88 (65% of overlapping DEGs) overlapping and concordant DEGs (**Fig 5B-C**). Among the 88 concordantly shared DEGs, we identified multiple expected angiogenic-related genes, including *Angpt2, Cxcr4, Esm1*, and *Etv4*. Surprisingly, we also identified shared decreased expression of retinoic acid receptor γ (*Rarγ*) and cytochrome P450 family 26 subfamily B member 1 (*Cyp26b1*), genes involved in and regulated by ATRA signaling, suggesting that decreased endothelial ATRA signaling may be a shared feature of AVM pathophysiology in both HHT and Glenn circulation (**Fig 5D**).

**Figure 5.**
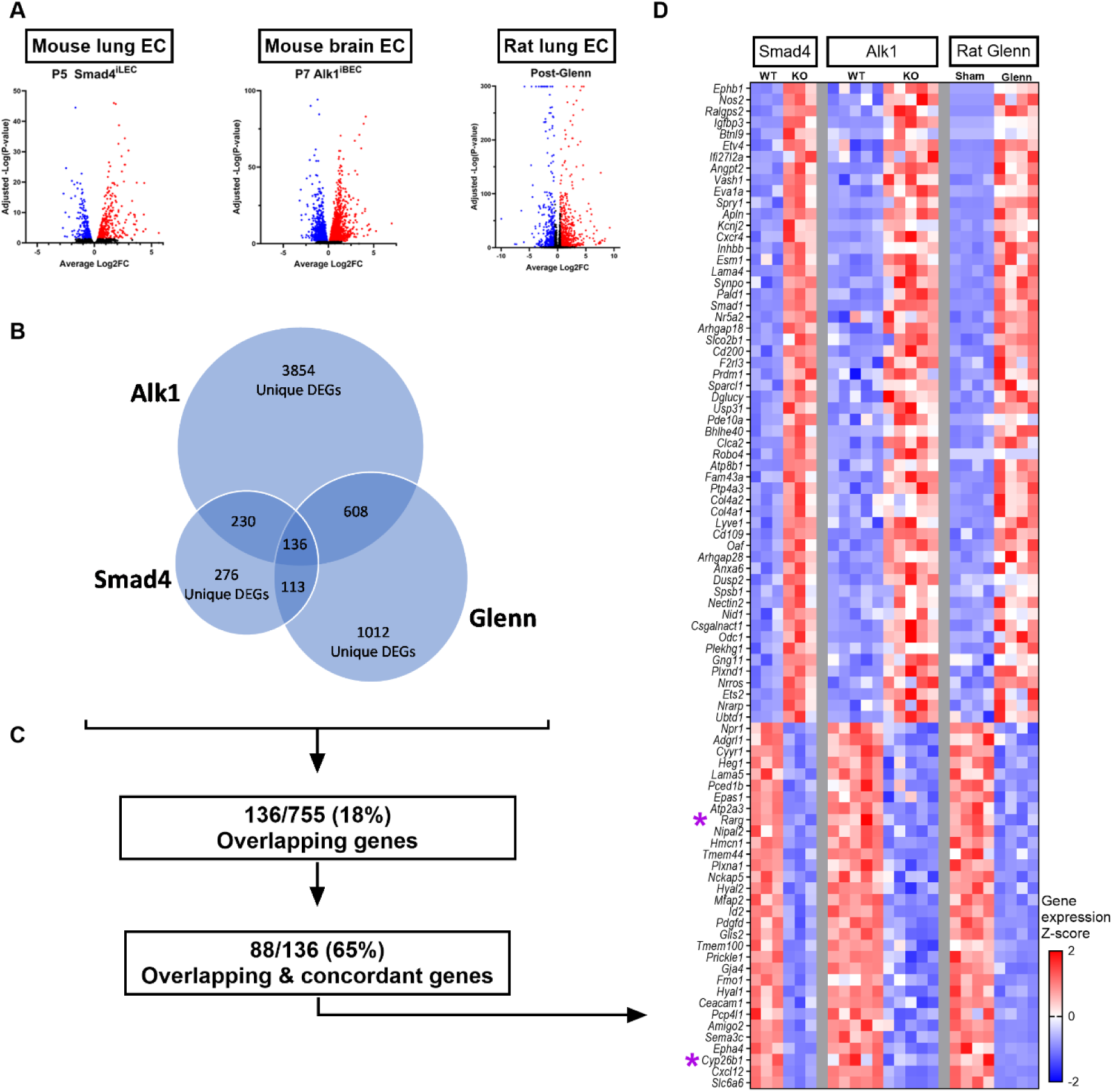
Conserved endothelial gene expression patterns in transgenic mouse models of hereditary hemorrhagic telangiectasia (HHT) and rat model of Glenn circulation. (A) Volcano plots for each model showing upregulated (red) and downregulated (blue) genes identified in endothelial-sorted bulk RNAseq (mouse models) and cluster 1 endothelial cells (ECs) in scRNAseq. DEG strictly defined based on adjusted p-value <0.05. (B) Venn diagram showing unique and overlapping DEGs among each model. (C) Flow chart specifying that there were 136 overlapping DEGs common to all 3 models (out of a possible 755 DEGs). Among the 136 overlapping DEGs, 88 (65%) were differentially expressed with log2FC in the same direction (i.e., concordant differential expression) in all 3 models. (D) Heatmap of all 3 models identifying the 88 overlapping and concordant genes. Purple asterisks identify genes involved in all-*trans* retinoic acid (ATRA) signaling (*RARγ* and *Cyp26b1*), indicating that decreased ATRA signaling is common in all 3 AVM models.

### Validation of decreased retinoid levels in post-Glenn lung endothelial cells

To determine the extent to which ATRA signaling may be dysregulated in lung ECs post-Glenn (**Fig 6A**), we further analyzed ATRA-related genes in our scRNAseq cluster 1 ECs and independently validated a subset of DEGs with CD31^+^ sorted qPCR (**Fig 6A-C**). In our scRNAseq cluster 1, we identified multiple DEGs associated with retinoid metabolism, including increased *Adh6a* (Log2FC= +1.2; Adj p-value = 1.7E-2; encodes a dehydrogenase enzyme that can oxidize ROH to retinaldehyde in the first step toward the active ATRA), increased *Nceh1* (Log2FC= +1.0; Adj p-value = 6.1E-8; encodes a putative RE hydrolase that converts stored RE to ROH) [44], decreased *Cyp26b1*(Log2FC= -3.2; Adj p-value = 1.6E-76; encodes a specific and potent catabolizing enzyme of ATRA), decreased *Dhrs3* (Log2FC= -1.1; Adj p-value = 4.0E-28; encodes a retinaldehyde reductase to limit excess levels of ATRA), decreased *Rarγ* (Log2FC= - 1.0; Adj p-value = 3.2E-9; encodes a *γ* isoform of retinoic acid nuclear receptor transcription factor that regulates ATRA-mediated gene expression), and decreased *Rarres1* (Log2FC= -2.9; Adj p-value = 3.1E-3; encodes a transmembrane protein involved in cell proliferation and whose expression is retinoic acid responsive) (**Fig 6A-B**). Independent validation of a subset of scRNAseq DEGs with CD31^+^ sorted qPCR confirmed decreased gene expression of *Cyp26b1* (p=0.02) and *Dhrs3* (p=0.02) (**Fig 6C**). Among the canonical retinoic acid nuclear receptor transcription factors, only *Rarγ* was decreased in Glenn rats (p=0.03) (**Fig 6C**).

**Figure 6.**
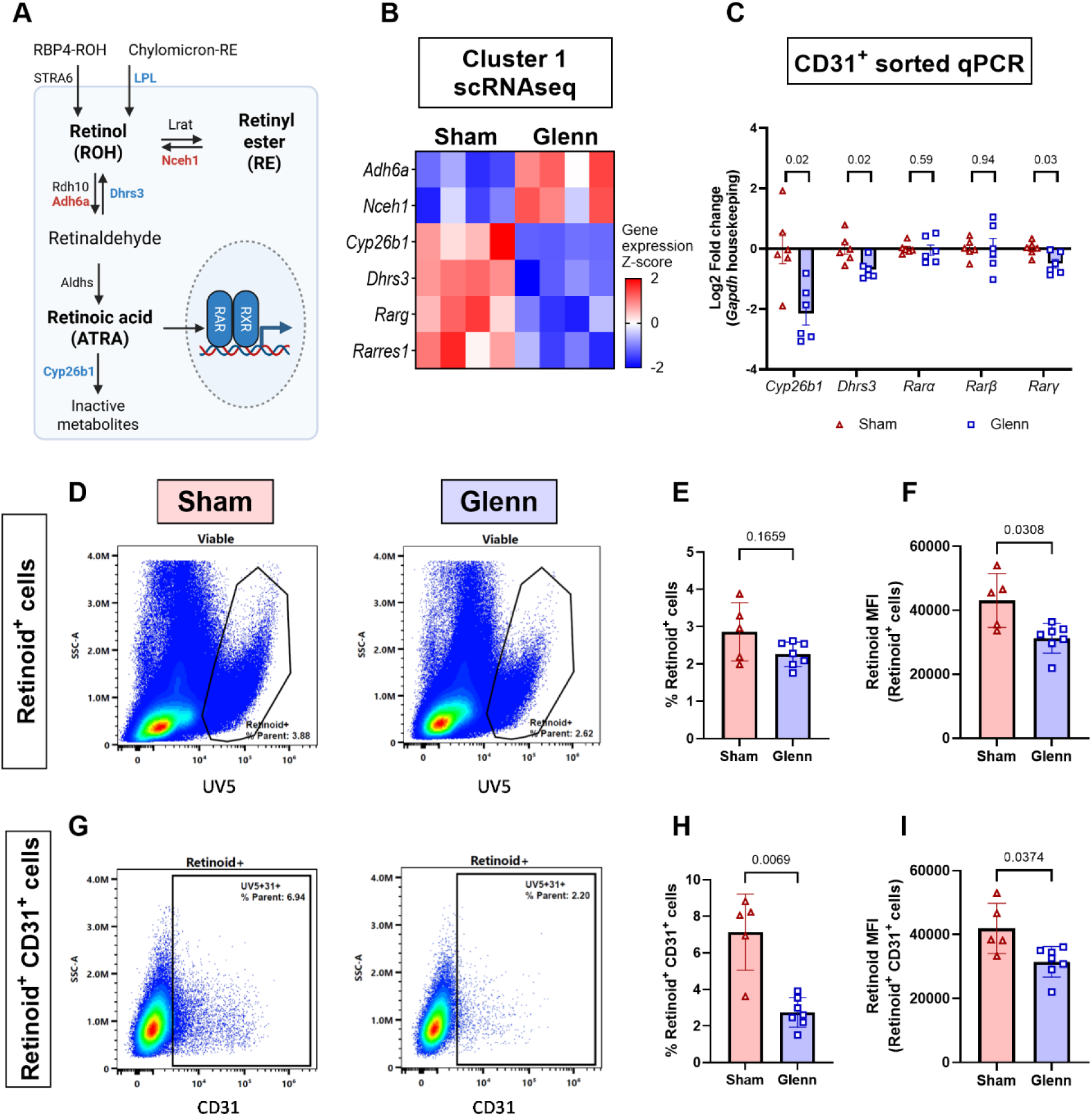
Decreased all-*trans* retinoic acid (ATRA) signaling and intracellular retinoid levels in lung endothelial cells (ECs) after Glenn. (A) Illustration of pathways for retinoid (retinol [ROH] and retinyl ester [RE]) uptake, metabolism, and signaling. Red colored gene names are upregulated genes in cluster 1 ECs, blue colored gene names are downregulated genes in cluster 1 ECs, and black colored gene names encode canonical enzymes in retinoid metabolism but are not detected (*Stra6*) or not differentially expressed (*Lrat, Rdh10, Aldh1a1*) in cluster 1 ECs. (B) Heatmap of gene expression z-score for significantly differentially expressed genes (DEGs) associated with ATRA signaling and metabolism in cluster 1 ECs. (C) qPCR validation of a subset of ATRA genes in CD31^+^ sorted lung cells after Glenn or sham surgery validating the scRNAseq data (n=6 sham and n=6 Glenn rats). (D-F) Spectral cytometry quantification of endogenous intracellular retinoid levels based on retinoid auto-fluorescence at UV5 spectral channel. There is no difference in the total amount of retinoid-containing (retinoid^+^ cells) in the lung, though there are decreased intracellular retinoid levels based on median fluorescent intensity (MFI) in post-Glenn rats (p=0.0308). Scatter plot with bars indicates mean ± SD (n=5 sham rats and n=7 Glenn rats). (G-I) Spectral cytometry quantification of retinoid^+^CD31^+^ lung cells identifies fewer retinoid^+^CD31^+^ cells after Glenn surgery (p=0.0069) and decreased retinoids levels (p=0.0374). Scatter plot with bars indicates mean ± SD (n=5 sham rats and n=7 Glenn rats).

Interestingly, we also identified significantly decreased expression of *Lpl* (scRNAseq cluster 1: Log2FC= -1.7; Adj p-value = 1.6E-76) (**Fig 3A**, **Fig 4A)** (CD31^+^ sorted qPCR: p<0.01) (**Fig 4C**), which encodes a lipase enzyme critical for RE hydrolysis from chylomicrons to facilitate lung EC-uptake of free ROH [40,45]. In contrast, *Stra6,* which encodes the cell surface receptor for ROH uptake of liver-derived ROH-RBP4 complex, was not detected in cluster 1 ECs nor via qPCR in CD31^+^ sorted cells. This is consistent with previously published data that lung ECs primarily uptake retinoids via chylomicron-derived retinoids and not liver-derived ROH-RBP4 [40].

Looking beyond gene expression, we further quantified endogenous retinoid levels in lung cells 3 weeks after Glenn or sham surgery using spectral cytometry based on retinoid autofluorescence. Among cells with strongly detectable intracellular retinoids, there was no difference in the percentage of retinoid^+^ cells (p=0.1659) (**Fig 6D-E**); however, there were significantly decreased levels of intracellular retinoids detected in these cells after Glenn (p=0.0308) (**Fig 6F**). Looking more specifically at lung ECs, there were significantly fewer retinoid^+^CD31^+^ lung cells (p=0.0069) (**Fig 6G-H**) and significantly decreased retinoid levels in these cells after Glenn (p=0.0374) (**Fig 6I**). Using HPLC to compare retinoid tissue concentrations in the lungs (ROH and RE), there were no statistical differences in whole lung Glenn samples compared to sham samples (**Supp Fig 6A-B**), which was similar to our spectral cytometry-based quantification of lung retinoids in all live lung cells (**Supp Fig 6C**). Collectively, these data suggest that retinoids are decreased specifically in lung ECs after Glenn, and retinoid levels are preserved in other non-EC cell types.

### Dietary vitamin A intake promotes progression of post-Glenn PAVMs

To investigate whether dietary intake of Vit A impacts PAVM pathogenesis, we first validated that dietary modification of Vit A intake decreases retinoid levels in the lung (**Supp Fig 7A-C; Supp Fig 8A-B**) and decreases retinoid levels specifically in lung ECs (**Supp Fig 7D-F**). As expected, HPLC quantification confirmed that dietary modification significantly impacted retinoid storage in the liver (**Supp Fig 8C-D**) but did not alter plasma ROH levels (**Supp Fig 8E**), supporting that circulating ROH levels do not reflect tissue-specific retinoid levels. In contrast, and as expected, dietary modification of Vit A significantly altered plasma RE levels (**Supp Fig 8F**) given that REs circulate in plasma in diet-derived chylomicrons.

After validating our dietary modification approach, we assessed whether modifying Vit A dietary intake impacts the functional effect of PAVM intrapulmonary shunting (**Fig 7A-B**). Similar to our previously published data at 2 weeks post-surgery, Glenn rats on Vit A control diet had increased shunting compared to the diet-matched sham control rats (p=0.0128) [15]. Significantly, Glenn rats on Vit A deficient diet had increased shunting compared to diet-matched sham control rats (p<0.0001), as well as compared to Glenn rats on Vit A control diet (p=0.0016) and Glenn rats on Vit A excess diet (p<0.0001). In fact, Glenn rats on Vit A deficient diet approached the progressive shunting we previously identified at 6 months post-Glenn on standard rat chow (1.54±0.23%) [15]. Equally important, Vit A deficient diet did not increase shunting in sham rats, while Glenn rats on Vit A excess diet did not exhibit increased shunting compared to diet-matched sham control rats (p=0.8243) (**Fig 7B**). Collectively, these data support that decreased retinoid availability may be critical for PAVM progression but not PAVM initiation, and that increasing lung EC retinoid availability may mitigate PAVM pathogenesis.

**Figure 7.**
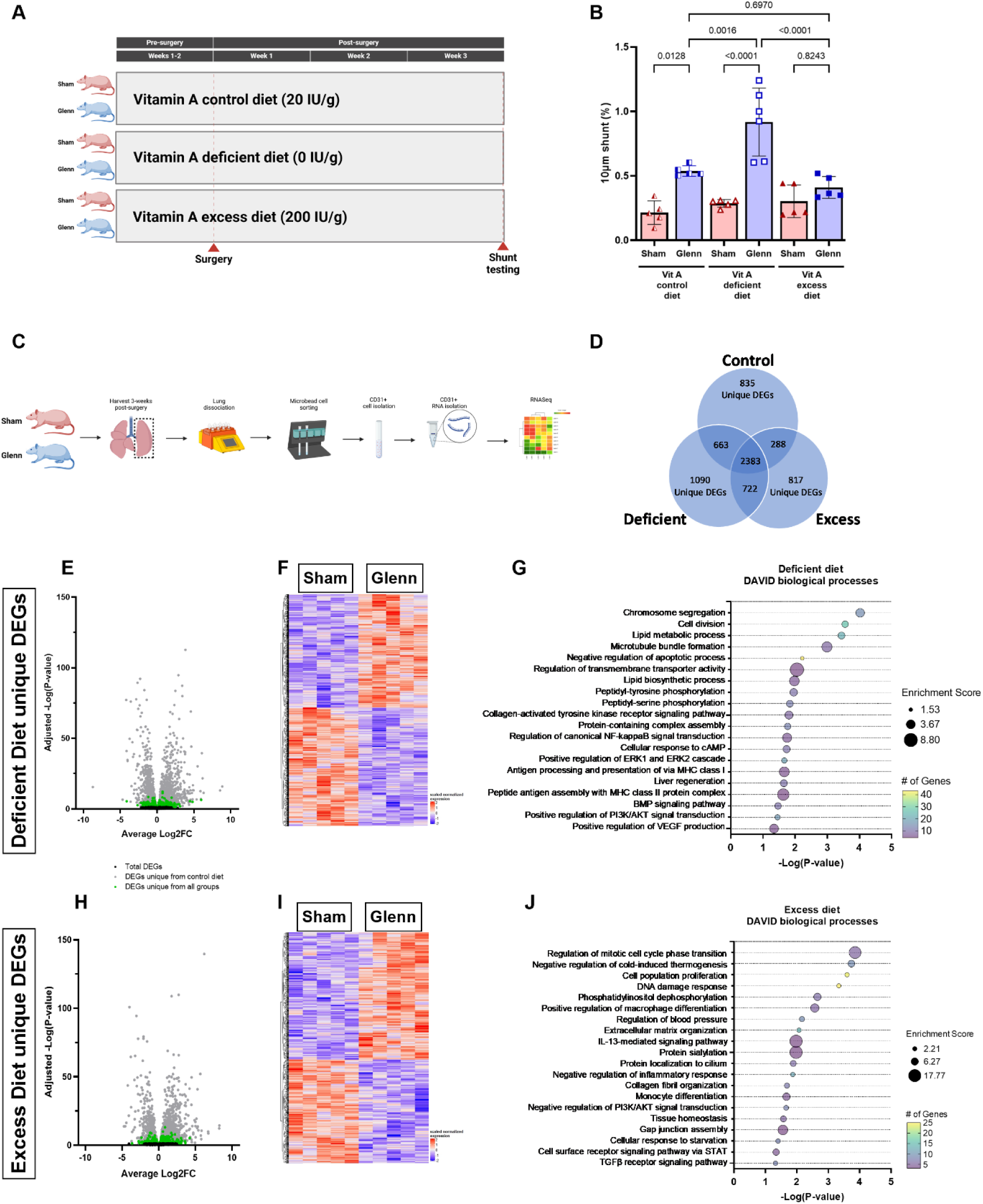
Decreased dietary vitamin A promotes progression of post-Glenn PAVMs. (A) Experimental schematic for modification of dietary vitamin A intake. Diets started 1-2 weeks prior to surgery at 6-7 weeks of age and continued for 3 weeks after surgery for assessment of CD31^+^-sorted RNAseq. (B) Quantification of intrapulmonary shunting with 10µm fluorescent microspheres identified increased shunting in Glenn rats on control diet compared to diet-matched sham control (p=0.0128), as well as increased shunting in Glenn rats on deficient diet compared to diet-matched sham control (p<0.0001) and Glenn rats on control diet (p=0.0016) and excess diet (p<0.0001). Glenn rats on excess diet did not have increased shunting compared to diet-matched sham control (p=0.8243) or any other group. Scatter plot with bars indicates mean ± SD (n=5-6 rats per group). (C) Experimental schematic for performing RNAseq on CD31^+^ sorted cells. (D) Venn diagram showing differentially expressed genes (DEGs) for each Glenn versus sham diet-matched comparison and the corresponding overlapping and unique DEGs for each group. (E-G) Volcano plot of DEGs, heatmap of unique DEGs, and impacted biological processes for the unique DEGs from rats on Vit A deficient diet. (H-J) Volcano plot of DEGs, heatmap of unique DEGs, and impacted biological processes for unique DEGs from rats on Vit A excess diet.

Next, after identifying that a Vit A deficient diet promotes post-Glenn PAVM progression, we sought to determine how dietary Vit A modification impacts EC gene expression by performing bulk RNAseq on sorted CD31^+^ lung cells (**Fig 7C**). After identifying DEGs (adjusted p-value <0.05) between Glenn and sham rats on each diet, we observed many reproducibly shared DEGs across all 3 diets (n=2383 shared DEGs) indicating a strong and reproducible Glenn-effect on lung ECs (**Fig 7D**). We also identified many unique DEGs specific to each diet, particularly with n=1090 unique DEGs on the Vit A deficient diet and n=817 unique DEGs on the Vit A excess diet (**Fig 7D**). Analyzing these unique DEGs with DAVID, we identified multiple significantly enriched biological processes (**Fig 7G, 7J**). Most notably, unique deficient diet DEGs were associated with positive regulation of PI3K/AKT signaling (p=3.5E-2), whereas unique excess diet DEGs were associated with negative regulation of PI3K/AKT signaling (p=2.2E-2) and phosphatidylinositol dephosphorylation (p=2.2E-3). Deficient diet DEGs were also associated with altered lipid metabolism (p=3.6E-4), lipid biosynthesis (p=1.1E-2), increased ERK1/2 signaling (p=2.2E-2), and others. Excess diet DEGs were additionally associated with multiple EC-non-autonomous biological processes, including macrophage differentiation (p=2.7E-3), monocyte differentiation (p=2.1E-3), and more. Altogether, these data suggest that decreased EC ATRA signaling may promote PAVM progression through increased PI3K signaling, whereas increasing EC ATRA may mitigate PAVM progression through decreased PI3K signaling and potentially EC-non-autonomous mechanisms involving crosstalk with innate immune cells.

## Discussion

This study is the first comprehensive analysis of how Glenn circulation alters lung gene expression at single-cell resolution early after Glenn surgery in a surgical rat model. With a modest number of biological replicates (n=4 per group), we identified significant transcriptional differences in lung ECs at this early stage of PAVM initiation. We identified that multiple molecular signaling pathways were dysregulated in ECs between Glenn and sham rats, including overlap with HHT pathophysiology, increased angiogenesis, and decreased ATRA signaling. Validation of our scRNAseq data confirmed decreased retinoid levels in lung ECs early after Glenn surgery. Importantly, we also showed that modification of dietary Vit A intake impacts lung EC gene expression and progression of post-Glenn PAVM shunting.

Previous groups reported their findings using animal models of unilateral Glenn circulation, though no studies have previously used next-generation sequencing, cell-specific sequencing, or scRNAseq [7–16]. More than 15 years ago, Tipps et al reported their findings after performing a microarray using total RNA from the whole left lung lysate in rats 8 months post-surgery [16]. They reported altered expression of genes associated with angiogenesis, vascular remodeling, vasodilation, and several other processes. Consistent with the seminal study by Tipps et al, many of these same expression patterns were observed in ECs in our scRNAseq experiment and confirmed with qPCR, including increased *Angpt2* and *Esm1*, and decreased *Acvrl1, Edn1, Glp1r, and Cyp26b1.* Moreover, we significantly expanded the scope and resolution of molecular profiling. We studied lung tissue early after surgery during the PAVM initiation phase, and we confirmed that these gene expression changes are specifically occurring in lung ECs.

Similar to the finding by Tipps et al of decreased *Acvrl1* expression, we identified differential expression of multiple genes associated with BMP-TGF-β signaling that overlap with the pathogenesis of HHT AVMs. Our scRNAseq dataset identified that two causative genes of HHT (*Acvrl1* and *Eng*) were downregulated in rat Glenn lung ECs, consistent with loss of function of these genes in HHT AVM pathogenesis. Additionally, we found multiple genes in this pathway were dysregulated in a similar manner to HHT AVM pathogenesis, including increased *Angpt2* and decreased *Id1, Id3,* and *Tmem100* [17,37,42–43,46]. This is consistent with a study by Bartoli et al previously reporting that increased angiopoietin signaling may contribute to single ventricle PAVMs [47]. Collectively, these data establish for the first time that there is overlap in the pathogenesis of single ventricle PAVMs and HHT AVMs. Accordingly, our comprehensive bioinformatic analysis using IPA predicts with a high statistical probability (p=7.5E-18) that GDF2 (also known as BMP9 and a known ligand for ALK1 and ENG) is a potential upstream regulator de-activated in Glenn circulation (activation Z-score -3.7). Capasso et al previously investigated patient blood samples to determine whether BMP9 might be enriched in hepatic vein blood; however, no differences in BMP9 plasma levels in hepatic vein blood vs SVC blood were found [48]. This suggests that non-pulsatile and decreased flow in Glenn circulation may have a critical role in single ventricle PAVM pathogenesis – potentially priming the vascular endothelium for disease or potentiating the effects of circulating biochemical factors in hepatic vein blood [49].

Beyond the conventional genes involved in BMP-TGF-β signaling, we identified conserved gene expression patterns in both HHT models and our Glenn model that are consistent with decreased ATRA signaling. Because ATRA signaling controls gene expression of hundreds of downstream genes and multiple downstream signaling pathways, decreased ATRA signaling may act as a central hub for coordinating vascular remodeling in AVM pathogenesis. Our dietary modification of Vit A intake significantly increased PAVM progression in our rat Glenn model, potentially by promoting PI3K signaling, which is a common driver of HHT-associated AVMs [46]; whereas, decreased Vit A intake did not initiate shunting in sham surgical rats. Collectively, these data suggest that ATRA signaling may act as a conserved and critical regulator of AVM pathogenesis such that decreased ATRA signaling promotes AVM progression. Whether ATRA can modify AVM pathophysiology in other vascular beds or AVM conditions remains unknown.

In addition to angiogenesis and ATRA signaling, we identified that lung ECs in Glenn circulation up-regulated cellular responses to potentially multiple inflammatory cytokines, including IL1, TNF, and LPS (**Figs 3-4**). To our knowledge, inflammation has not been previously identified or proposed as a pathologic trigger for single ventricle PAVMs, although inflammation is a well-recognized aspect of HHT-related AVMs [50–55]. Decreased lung EC ATRA signaling specifically in lung ECs is associated with increased lung leukocyte infiltration even without exogenous inflammatory stimuli [40], suggesting that ATRA signaling may in part be contributing to post-Glenn inflammation. Additionally, we previously identified that SVC serum obtained from patients with Glenn circulation was enriched in multiple cytokines and chemokines [56]. Whether broad or specific anti-inflammatory therapies can modify single ventricle PAVM pathogenesis, or whether ATRA signaling is contributing to post-Glenn inflammation and potentially leukocyte recruitment, is currently untested and unknown but warrants further investigation.

Lastly, we identified gene expression changes in lung ECs associated with vasodilation, specifically down-regulation of a potent vasoconstrictor (*Edn1*) and up-regulation of a potent vasodilator (*Nos2*). The stimuli for these changes are unknown; however, early vasodilation could be an initial physiologic cause of intrapulmonary shunting and the PAVM phenotype. We speculate that this early vasodilatory response is the initial manifestation of single ventricle PAVM shunting and sustained vasodilation combined with pro-angiogenic and pro-inflammatory signaling ultimately leads to architectural remodeling of the lung vasculature if hepatic vein blood perfusion is not restored. The specific factor(s) in hepatic vein blood remain unknown, but we are poised to identify causative factor(s) with our current animal model.

Our study has several important limitations. First, despite validation with independent methods, gene expression changes identified in this study may be related to tissue remodeling post-Glenn but indirectly or even independently of PAVM pathogenesis post-Glenn. For example, formation of aortopulmonary collaterals is a universal phenomenon in patients post-Glenn [8, 57], so we anticipate that aortopulmonary collaterals similarly develop in our post-Glenn rats and may cause gene expression changes independent of PAVMs. Second, in contrast to patients with single ventricle circulation and systemic hypoxemia, our rats with Glenn and sham surgeries do not experience significant hypoxemia [15]. It is possible that systemic hypoxemia may contribute to relevant gene expression differences that our rat model does not fully recapitulate.

In summary, we report the novel application of scRNAseq to study single ventricle PAVMs in a surgical rat model of Glenn circulation. Using a comprehensive and unbiased approach, we identified that ECs are the primarily affected cell type in the lungs early post-Glenn, and multiple biological processes are dysregulated in rat lung ECs post-Glenn, including decreased ATRA signaling and conserved gene expression patterns with HHT mouse models. Dietary modification of Vit A intake altered post-Glenn shunting and represents a novel potential therapeutic strategy for single ventricle PAVMs and HHT AVMs.

## Acknowledgements

This study was supported by the National Institutes of Health from the National Heart, Lung, and Blood Institute (K08HL157510 – ADS; R01HL171112 – IS, R33HL154254 and R01HL179583 – RR, R01HL163196 and R01HL139713 – SMM), Department of Defense Investigator-Initiated Research Award (PR160198 – SMM), Medical College of Wisconsin Department of Pediatrics (ADS), Herma Heart Institute Innovation Funds (ADS), American Heart Association Predoctoral Fellowship (20PRE35120237 – XZ), and joint CTSI Pilot – Mellowes Center for Genomic Sciences and Precision Medicine award (UL1TR001436 – ADS).

## Study Approval

Experimental rat protocols were approved by the Medical College of Wisconsin Institutional Animal Care and Use Committee prior to initiation of experimental protocols (Animal Use Agreement #7731). Experimental mouse procedures were approved by Tulane University’s institutional Animal Care and Use Committee policies and adhere to ethical standards.

