## Supplemental Material for "Decreased endothelial cell retinoic acid signaling accelerates progression of single ventricle pulmonary arteriovenous malformations"

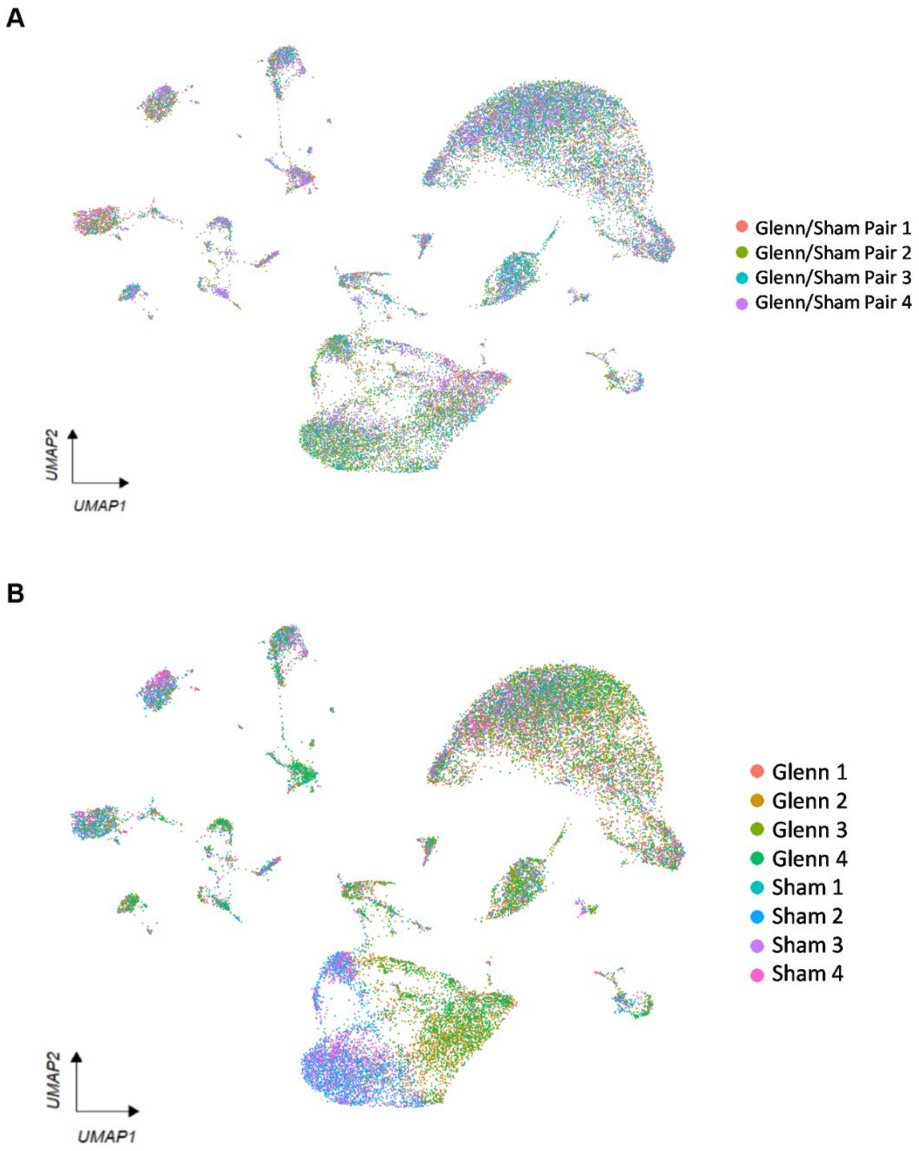


**Supplemental Figure 1. Source of lung cells within the global un**iform manifold approximation and projection (UMAP) plot. (A) UMAP plot color coded by Glenn and sham pairs that had lung tissue harvested simultaneously demonstrates relatively equal distribution of harvest pairs in all clusters. (B) UMAP plot color coded by individual animals demonstrates separation of Glenn and sham samples within clusters 1 and 3.

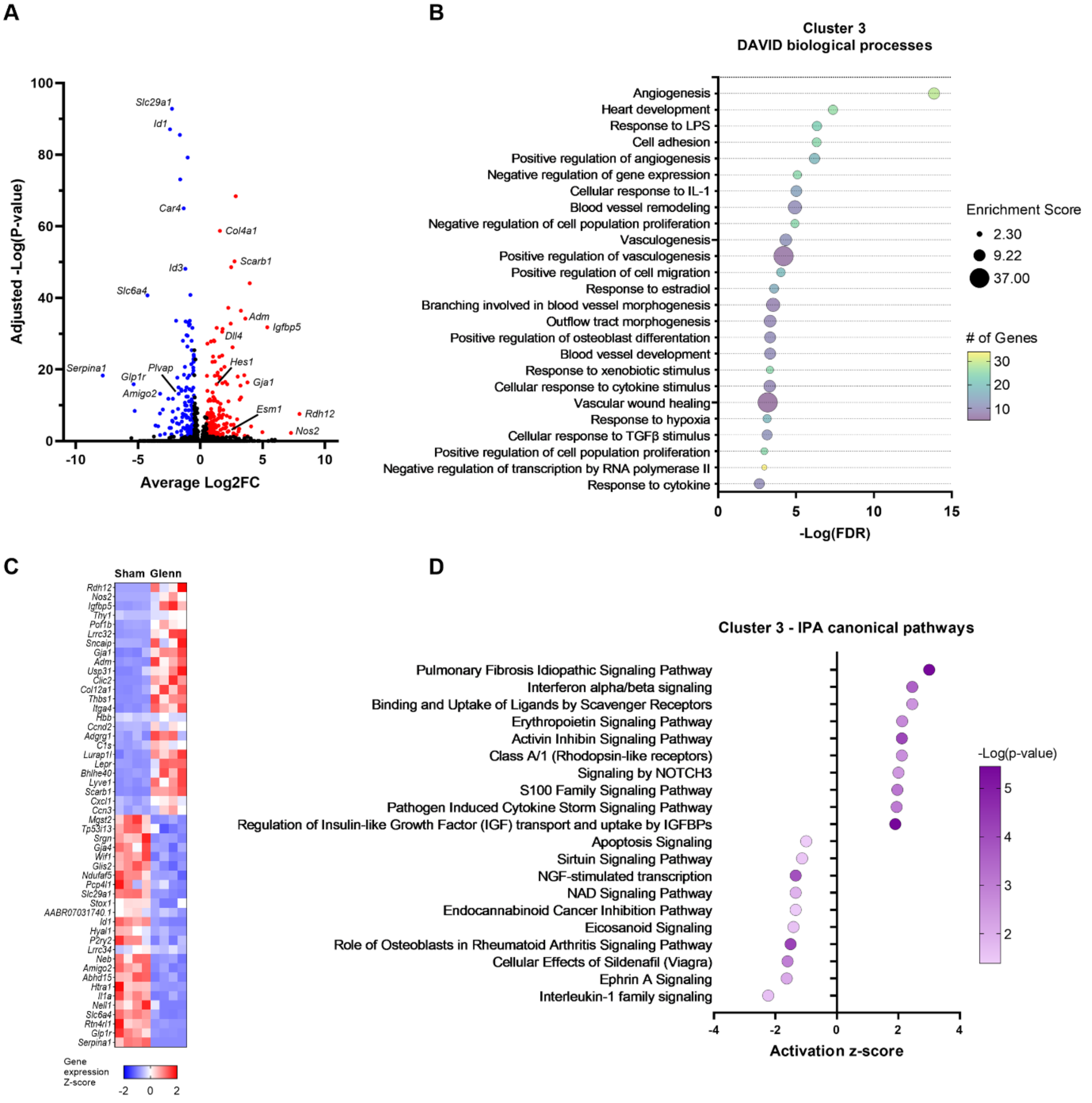


**Supplemental Figure 2. Capillary lung endothelial cells (ECs) comprise cluster 3 cells and are significantly dysregulated in Glenn versus sham rats.** (A) Volcano plot showing upregulated (red) and downregulated (blue) genes identified in cluster 3 with annotation of notable differentially expressed genes (DEGs). (B) Dot plot showing the top 25 biological pathways identified using DAVID pathway analysis and cluster 3 DEGs. (C) Heatmap of the top 25 most up-regulated and top 25 most down-regulated DEGs in cluster 3. Each column represents an individual rat, and each row represents a gene. (D) Dot plot of the 10 most activated and 10 most de-activated canonical pathways identified using Ingenuity Pathway Analysis (IPA) and the entire cluster 3 dataset. Positive activation z-score indicates increased activity of the associated pathway in Glenn rats, whereas negative activation z-score indicates decreased activity of the associated pathway in Glenn rats.

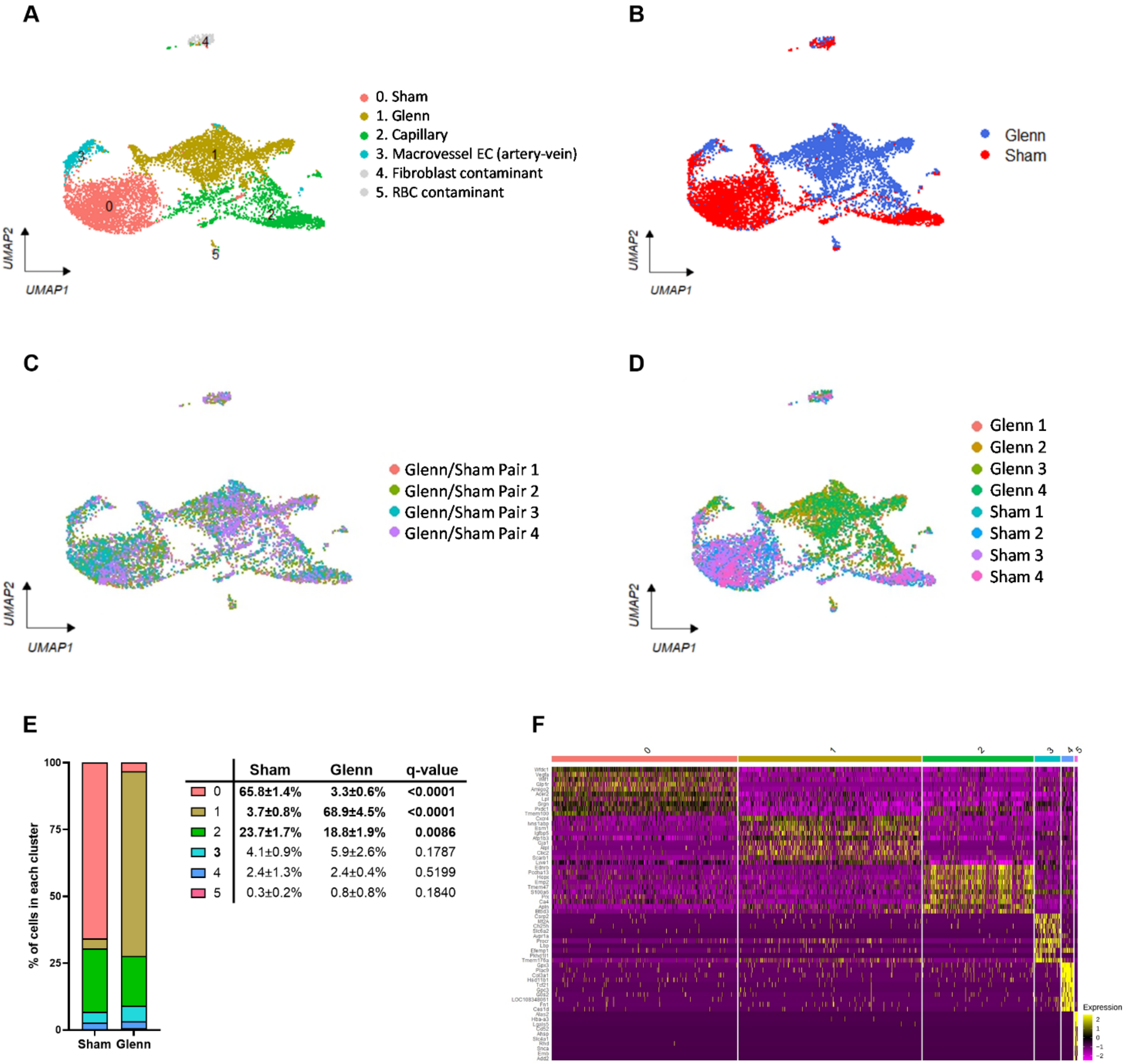


**Supplemental Figure 3. Surgical group is the strongest factor controlling endothelial cell (EC) sub-clustering.** (A) Combined uniform manifold approximation and projection (UMAP) plot with all samples (n=8) showing EC sub-clusters identified with unsupervised clustering and subsequent cluster annotation. Sub-clusters 0 and 1 are annotated by surgical group because differentially expressed genes (DEGs) between the two surgical groups were the primary distinguishing features of these sub-clusters as opposed to canonical EC markers of arteriovenous specification. (B) UMAP plot color coded by surgical group (Glenn versus sham) demonstrates pronounced separation within sub-clusters 0 and 1 between Glenn and sham rats. (C) UMAP plot color coded by Glenn and sham pairs that had lung tissue harvested simultaneously demonstrates relatively equal distribution of harvest pairs in all sub-clusters. (D) UMAP plot color coded by individual animals demonstrates separation of Glenn and sham samples between clusters 0 and 1. (E) Stacked bar graph with accompanying table indicates the mean ± SD of percent cells within each sub-cluster from each rat. Q-value indicates the adjusted p-value for multiple unpaired t-tests comparing the frequency of cells within each cluster. ECs within sub-clusters 0 and 1 were nearly absent in Glenn and sham rats, respectively. Sub-cluster 2 capillary ECs were significantly decreased in Glenn rats compared to sham rats, similar to findings within cluster 3 of the entire lung analysis (see Figure 2). (F) Global heatmap of all sub-clusters showing transcriptional gene expression differences for the top 10 genes within each cluster. Each column represents an individual cell and each row represents a gene.

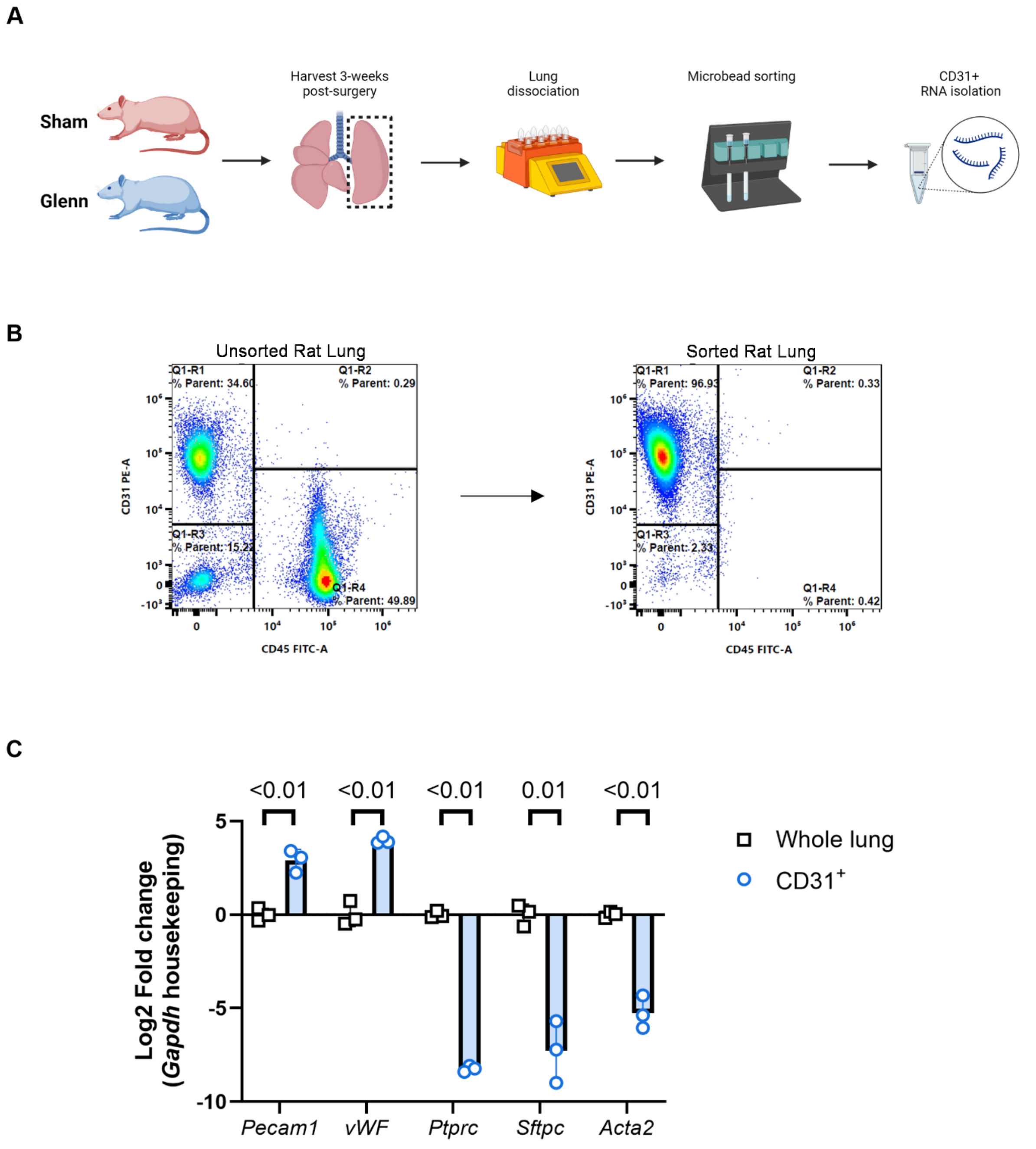


**Supplemental Figure 4. Validation of magnetic microbead methodology for sorting rat lung cells to isolate high purity of CD31$^{+}$ lung endothelial cells (ECs).** (A) Schematic for experimental methodology for dissociating rat lung tissue, magnetic microbead sorting lung cells, and isolating RNA from CD31$^{+}$ lung cells from rats. (B) Flow cytometry scatter plots validating successful magnetic microbead sorting of $CD31^+$ cells from dissociated rat lung tissue. The upper left quadrant for each scatter plot represents $CD31^+$ cells. (C) Scatter plot with bars (mean ± SD) shows validation of specific gene targets using qPCR from unoperated rats (whole lung n=3, $CD31^+$ sorted n=3). *Pecam1* and *vWF* (von Willebrand Factor) represent endothelial-specific genes. *Ptprc* (CD45), *Sftpc* (surfactant protein c), and *Acta2* (smooth muscle actin) represent genes enriched in leukocytes, lung epithelial cells, and mural cells, respectively.

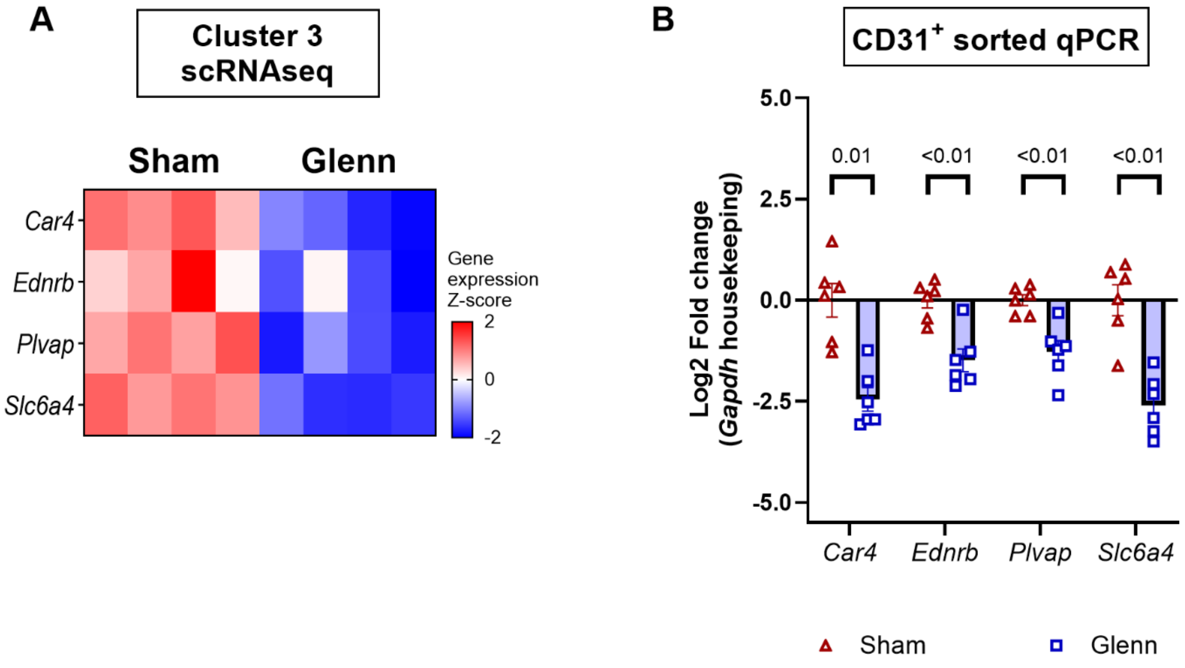


**Supplemental Figure 5. Glenn circulation down-regulates gene expression of multiple lung capillary markers**. (A) Heatmap of gene expression z-scores for differentially expressed genes (DEGs) in cluster 3 capillary endothelial cells. (B) Scatter plot with bars (mean ± SD) shows validation of capillary markers using quantitative RT-PCR (qPCR) (n=6 sham and n=6 Glenn rats).

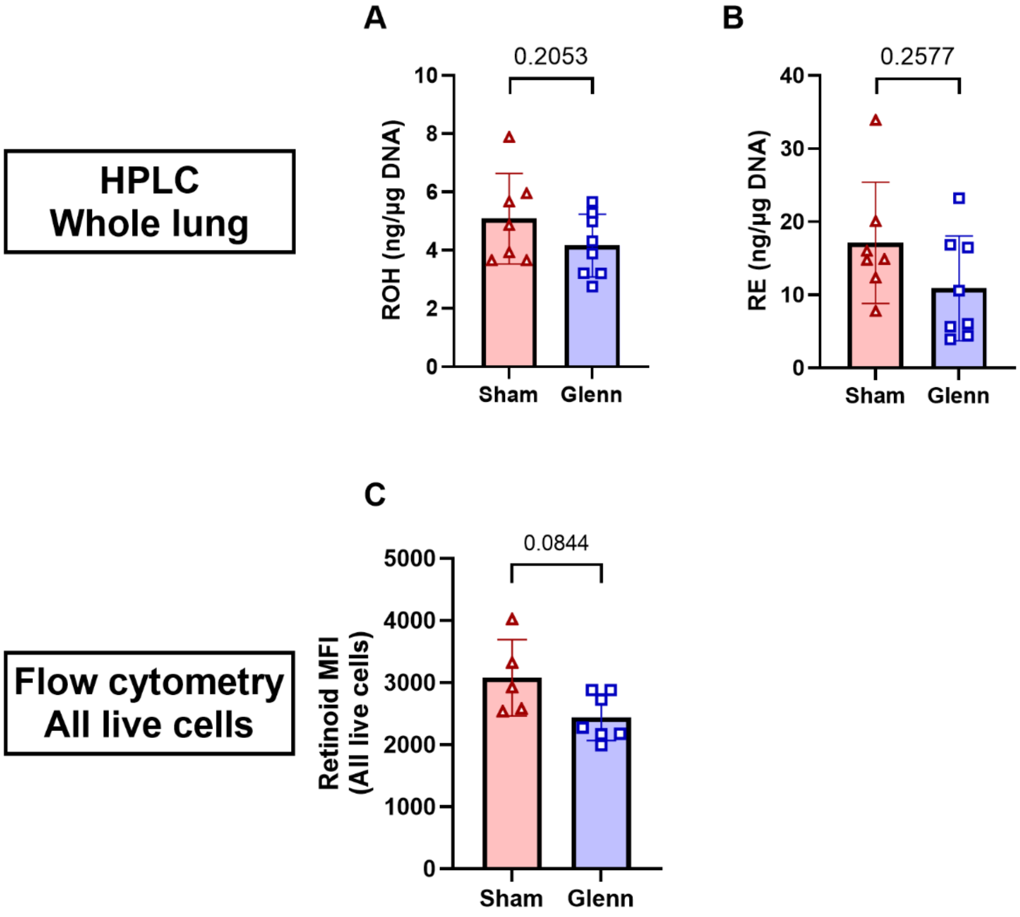


**Supplemental Figure 6. No difference in retinoid levels when analyzing the whole lung without sub-analyzing cell types**. (A-B) Quantification of whole lung retinoid levels (retinol [ROH] and retinyl esters [RE]) by high-performance liquid chromatography normalized to DNA content to account for cell number variability. (C) Quantification of whole lung retinoid levels by spectral cytometry based on endogenous retinoid auto-fluorescence at UV5 spectral channel (excitation 350nm, emission 455nm).

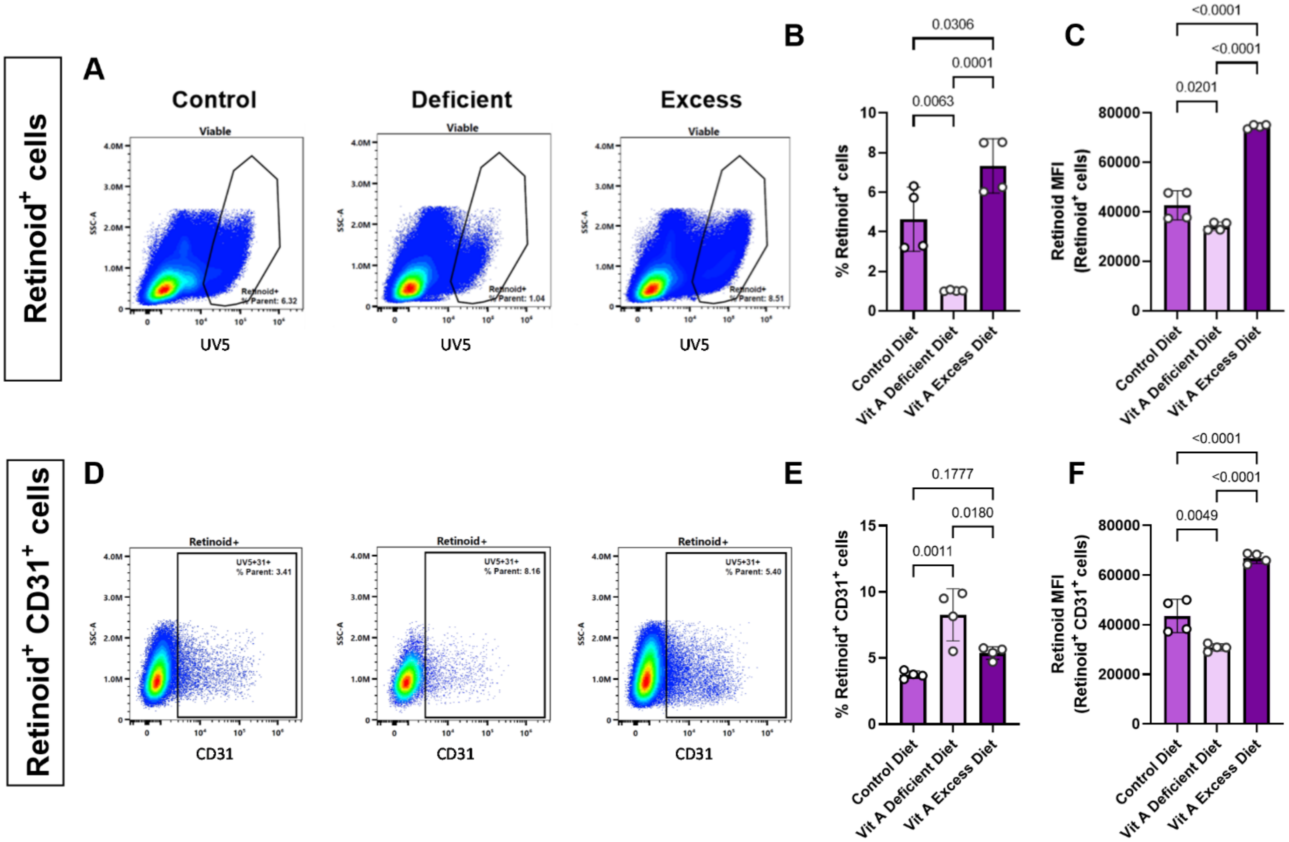


**Supplemental Figure 7**. **Dietary modification of Vitamin A intake alters retinoid levels in lung endothelial cells.** (A) Spectral cytometry quantification of endogenous retinoid storage based on retinoid auto-fluorescence. (B) There is decreased number of retinoid$^{+}$ lung cells in rats on Vit A deficient diet and increased number of retinoid$^{+}$ lung cells in rats on Vit A excess diet. (C) There are also decreased amounts of retinoids stored in retinoid$^{+}$ lung cells in rats on Vit A deficient diet based on median fluorescent intensity (MFI) and increased amount of retinoids stored in retinoid$^{+}$ lung cells in rats on Vit A excess diet. (D-F) Spectral cytometry quantification of retinoid$^{+}$ CD31$^{+}$ lung cells with percent of retinoid$^{+}$ CD31$^{+}$ lung cells (of all retinoid$^{+}$ lung cells) and retinoid MFI in retinoid$^{+}$ CD31$^{+}$ lung cells. There are significantly decreased amounts of retinoids stored in retinoid$^{+}$ CD31$^{+}$ lung cells for rats on Vit A deficient diet.

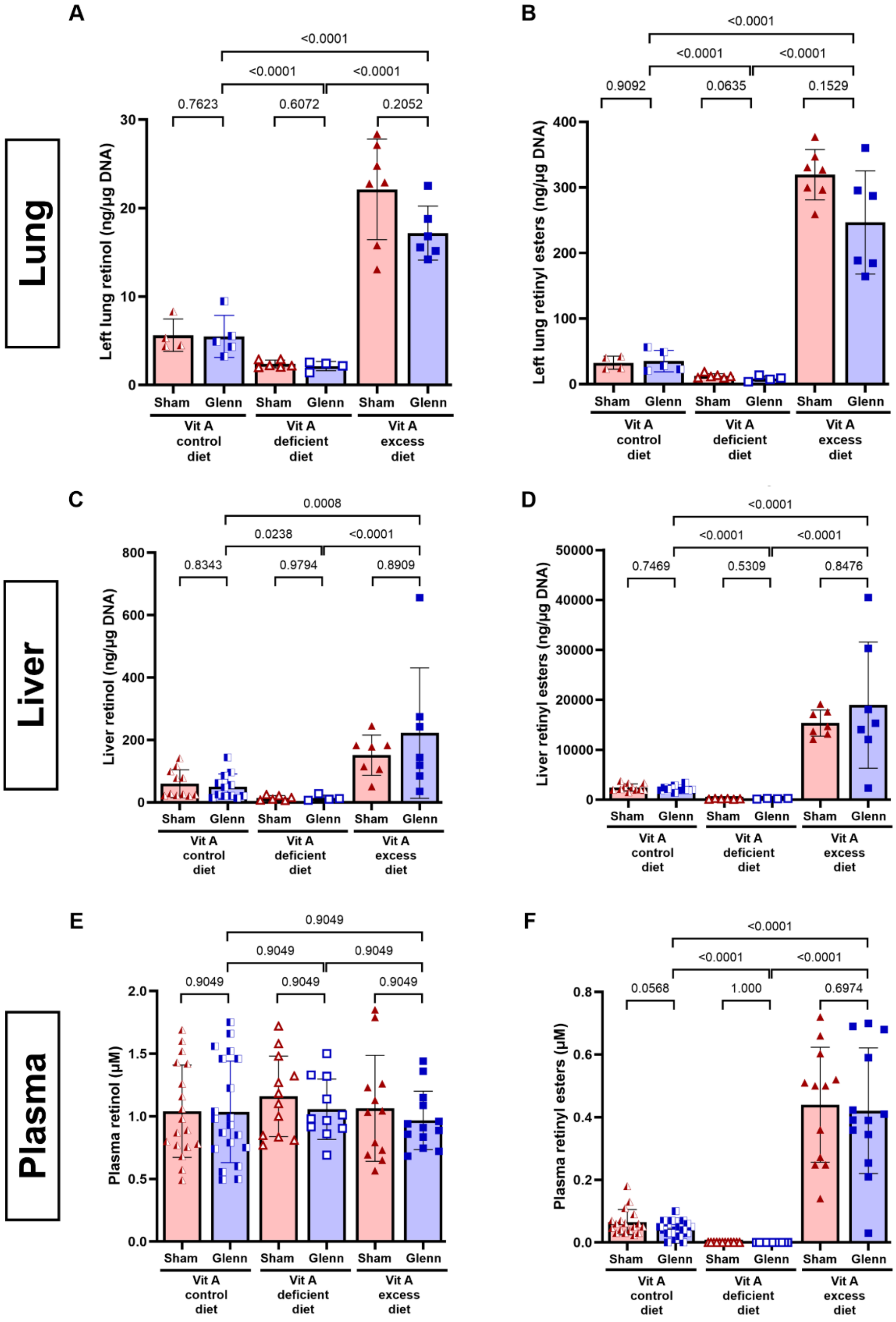


**Supplemental Figure 8. Retinoid levels in lung, liver, and plasma after dietary modification of Vitamin A intake.** (A-B) Quantification of whole lung retinoid levels (retinol [ROH] and retinyl esters [RE]) by high-performance liquid chromatography (HPLC) normalized to DNA content (n=4-7 per group). (C-D) Quantification of whole liver retinoid levels (ROH and RE) by HPLC normalized to DNA content (n=4-12 per group). (E-F). Quantification of plasma retinoid levels (ROH and RE) by HPLC (n=11-19 per group). Data were log-transformed prior to statistical analysis. P-values represent FDR-adjusted p-values for multiple comparisons.

| Gene Target | Forward Primer (5’-3’) | Reverse primer (5’-3’) |
|---|---|---|
| *Acta2* | AGCTACGAACTGCCTGATGG | TAGGTGGTTTCGTGGATGCC |
| *Acvrl1* | AAAGGAAATTCGGCCCCTGT | CCAAGTTCTCCCAAGGTGCA |
| *Adm* | TTTCAGCAGGGTATCGGAGC | TTCGGAACTGCGAGGAAGTG |
| *Angpt2* | AATGCACAGTAGCCCCTTCC | ATTTCTCCAGACCCGCAGTG |
| *Apoe* | GTGAGCAGATGGAGGAGGTG | TGCATGTCTTCCACTAGCGG |
| *Car4* | AGATTCAAGCCAAGGAGCCC | GTGAAGGGTGTCAGACTGGG |
| *Cd34* | GGCCATTCAGCAAGACAACA | TGAAGGCAGCATCAGTCAGA |
| *Cxcr4* | CGTGGGGATCAGCATCGATT | TAGAGGATGGGGTTCAGGCA |
| *Cyp26b1* | AGATGACACCGGCTTTGAGG | TGTAGGCTCCTGTACCCCAA |
| *Dhrs3* | GCAACAAAACCAGGCCCTTC | ACAGGTGTAAGTCCCCGAGA |
| *Edn1* | TGCCTCTTCTTGCTGTCTGG | CTACAGAAACTCCGCCCTGC |
| *Ednrb* | GAGTCCCGCCAAGATCCTTC | AGTTCCCGATGATGCCTAGC |
| *Esm1* | CAAGTATGCGGTGGATTGCC | GTACGGTAGCAGGTTTCCCC |
| *Foxf1* | TGTGTGATGTGAGGTGAGGC | TGGCCCTTTTCTCTTGCTGA |
| *Gapdh*[*] | TGTGAACGGATTTGGCCGTA | GATGGTGATGGGTTTCCCGT |
| *Gja1* | ACGTGGAGATGCACCTGAAG | CCACTGGATGAGCAGGAAGG |
| *Glp1r* | GGGCTTTATGGTGGCTGTCT | TTCATGCTGCTGTCCCTCTG |
| *Hspg2* | CCATACATCGACGCTTCCCA | TGGTTGAATGGGGTTGCCTT |
| *Il6st* | CTGCCCAGATTGTGTCACCT | TCGATACCAACGGCACCAAA |
| *Lpl* | ATGGACGGTGACAGGAATGT | CTGGATAATGTTGCTGGGCC |
| *Nos2* | ATTCCCAGCCCAACAACACA | GGTCGATGGAGTCACATGCA |
| *Pde2a* | CTGCTTCCACTACACAGGCA | CCAGGTGGGTGAAGAGGTTC |
| *Pecam1* | CGTCATTCCTCAGTCTCGGG | TTTGTCCACGGTCACCTCAG |
| *Pfkfb3* | CGCCGAATACAGCAACGAAG | GAGCCCCACCATCACAATCA |
| *Pla2g4a* | CCCTTGATTCTGCGACCTCA | TAGCCCACTTCTCTGCAAGC |
| *Plvap* | CATCGCCGCTATCATCCTGA | TGACCACCTCCATCTCCAGT |
| *Ptprc* | TCGGCCCAGAAGTCTTTGTC | GCTTCGTTTGAGGGGTAGCT |
| *Serpina1* | TTCAGCCTATACCGGGAGCT | TGTTTGCGAGTGTCACCCTT |
| *Sftpc* | GTCCTTGTCGTCGTGGTGAT | AGCGATGGTGTCTGTGTGTT |
| *Rarα* | GACTTGGTCTTTGCCTTCGC | ATGTCCACCTTGTCTGGCTG |
| *Rarβ* | GGGAGAACTTGGGATCGGTG | CCCAGCCCCGAATCATGAAT |
| *Rarγ* | CGGCTGCAAAAGTGTTTCGA | GCTGACCTTGGTGATGAGCT |
| *Scarb1* | GGGCAAACAGGGAAGATCGA | GTGAAGAACCTGGGGCATCA |
| *Slc1a5* | GTCCTGTACGGTCCTCAACG | CACGTCGTTCTTCACCTGGA |
| *Slc6a4* | GCATACGTGGTGACTCTGCT | AAGCCCAGCATCTCCTTCAC |
| *Sparcl1* | TGGAGACCACCCCATTGAAC | GTGTCAAGATCCTGTCCGCA |
| *Tgfbr1* | GGTCCAGTCTGCTTCGTCTG | GGTGGTGCCCTCTGAAATGA |
| *Tgfbr2* | TCACTAGGCACGTCATCAGC | GAACAATGGGCATCTTGGGC |
| *Tmem100* | AGACATTTCACCCTACCGCC | TTCGGTCCTCTCTATGCCCT |
| *Vegfa* | TCTTCAAGCCGTCCTGTGTG | GCTGGCTTTGGTGAGGTTTG |
| *vWF* | GACCCCTACGACTTTGCCAA | CGACGCCGTCTTCAGTAACT |

**Supplemental Table 1. Quantitative RT-PCR Primer sequences.**

* Reference #34 (Wan et al, 2023).

| Vitamin A Control diet | | Vitamin A Deficient diet | | Vitamin A Excess diet | |
|---|---|---|---|---|---|
| Component | g/kg | Component | g/kg | Component | g/kg |
| Casein (vitamin-free) | 193.0 | Casein (vitamin-free) | 193.0 | Casein (vitamin-free) | 193.0 |
| DL-Methionine | 3.0 | DL-Methionine | 3.0 | DL-Methionine | 3.0 |
| Sucrose | 504.3 | Sucrose | 505.7 | Sucrose | 505.3 |
| Corn starch | 150.0 | Corn starch | 150.0 | Corn starch | 150.0 |
| Cottonseed oil | 50.0 | Cottonseed oil | 50.0 | Cottonseed oil | 50.0 |
| Cellulose | 50.0 | Cellulose | 50.0 | Cellulose | 50.0 |
| Mineral Mix, AIN-76 | 35.0 | Mineral Mix, AIN-76 | 35.0 | Mineral Mix, AIN-76 | 35.0 |
| Calcium carbonate | 4.0 | Calcium carbonate | 4.0 | Calcium carbonate | 4.0 |
| Vitamin Mix, Teklad | 10.0 | Vitamin Mix, without choline, A, D, E | 5.0 | Vitamin Mix, without choline, A, D, E | 5.0 |
| Blue food color | 0.1 | Choline dihydrogen citrate | 3.497 | Choline dihydrogen citrate | 3.497 |
| Aspirin | 0.61 | Vitamin D3, cholecalciferol (500000 IU/g) | 0.0044 | Vitamin D3, cholecalciferol (500000 IU/g) | 0.0044 |
| | | Vitamin E, DL-alpha tocopheryl acetate (1000 IU/g) | 0.121 | Vitamin E, DL-alpha tocopheryl acetate (1000 IU/g) | 0.121 |
| | | Aspirin | 0.61 | Vitamin A palmitate (500000 IU/g) | 0.4 |
| | | | | Orange food color | 0.1 |
| | | | | Aspirin | 0.61 |

**Supplemental Table 2. Custom rat diets from Teklad Diets (Inotiv).**
